# Genomic structure and diversity of oak populations in British Parklands

**DOI:** 10.1101/2021.03.05.434114

**Authors:** Gabriele Nocchi, Nathan Brown, Tim Coker, William Plumb, Jonathan Stocks, Sandra Denman, Richard Buggs

**Author notes:** **Corresponding authors:** Gabriele Nocchi, Richard Buggs.

## Abstract

The two predominant oak species in Britain are *Quercus robur* (English or pedunculate oak) and *Q. petraea* (sessile oak). We sequenced the whole genomes of 386 oak trees from four British parkland sites and found over 50 million nuclear single nucleotide polymorphisms (SNPs), allowing us to identify 360 *Q. robur*, ten *Q. petraea* and 16 hybrid individuals using clustering methods. Comparing *Q. robur* and *Q. petraea* trees from Attingham Park, we found that the nuclear genomes of the two species are largely undifferentiated but identified 81 coding regions exhibiting strong interspecific differentiation. The nuclear genomes of our 360 *Q. robur* individuals showed no clear differentiation among the four parkland sites. Scans for selective sweeps in *Q. robur* highlighted regions containing genes with putative involvement in stress tolerance, one of which was moderately differentiated from *Q. petraea*. Reconstructions of past effective population sizes suggested a long population size decline in both *Q. robur* and *Q. petraea* over the Pleistocene, but population growth after the last glacial maximum. We assembled the whole chloroplast genomes of 287 *Q. robur*, 8 *Q. petraea* and 14 hybrid trees. In a phylogenetic network, these fell into five major haplotypes, which were shared among species but differed in frequency among parkland sites. We matched our chloroplast genome haplotypes to restriction enzyme fragment haplotypes identified in older studies that had surveyed ancient woodlands in Britain and much of Europe. This suggested that the parkland populations in our study derive from local seed sources.

## Introduction

Oaks (genus *Quercus*) are some of the most common and widely distributed forest trees in the northern hemisphere, found throughout North America, Europe and Asia comprising between 300 and 500 species (Kremer et al., 2012; Plomion and Fievet, 2013), with 435 included in the latest classification (Denk et al., 2017). Differentiating oak species morphologically has been termed a “botanical nightmare” due to ambiguous and extremely variable phenotypes, which is partly due to extensive hybridization and backcrossing (Denk et al., 2017; Kleinschmit, 1993; Kremer et al., 2012; Lepais et al., 2009; Leroy et al., 2019). A recent phylogenomic study showed that the oak phylogeny is highly reticulate, with phylogenetic incongruence widespread throughout the genome (Hipp et al., 2019).

The Eurasian white oaks form a clade in the phylogeny of Hipp et al. (2019) that is nested within clades of largely American oak species. White oaks are thought to have crossed the North Atlantic land bridge in the Oligocene and split into European and Asian clades in the Miocene, which then rapidly diversified (Hipp et al., 2019). Evidence from chloroplast variation and fossil pollen suggest that in the last glacial maximum (~20 Kya), European white oaks were constrained to refugia in southern Europe (Dumolin-Lapegue et al., 1997; Petit et al., 2002a; Petit et al., 2002b). Six chloroplast lineages have been identified corresponding to glacial refugia in the Balkans (lineage A), in western Spain (lineage B), in the south of Italy (lineage C), in eastern Spain (lineage D), in the eastern Balkans (lineage E) and north-east of the Black Sea (Crimea) (lineage F) (Petit et al., 2002a; Petit et al., 2002b). The vast majority of Britain’s native oak trees have been identified as belonging to lineage B (Cottrell et al., 2002).

In Europe today, the two most common oak species are *Q. robur* (English oak or pedunculate oak) and *Q. petraea* (sessile oak). These species are found from central Spain to the Urals and from Scandinavia to the south of Italy (Eaton et al., 2016). Their lineages seem to have diverged in the late Miocene/early Pliocene, and they are found in sister clades that each also contain a small number of species distributed in southern Europe and/or North Africa (Hipp et al., 2019).

*Quercus robur* and *Q. petraea* are sympatric across most of their geographic range but are morphologically distinct (Jones, 1959) and exhibit different ecological preferences. *Q. robur* reaches more northerly and easterly ranges in Europe, prefers moist and alkaline soils with high nutrient availability and it is more tolerant to periodic flooding and water logging (Barreneche et al., 1998; Eaton et al., 2016; Saintagne et al., 2004). Contrarily, *Q. petraea* prefers drier and more acidic soils and it is more resistant to drought (Barreneche et al., 1998; Eaton et al., 2016; Saintagne et al., 2004). *Q. robur* is a pioneer species able to colonize open ecosystems while *Q. petraea* appears to be more of a successional species that expands in area already occupied by *Q. robur* but can also behave as a pioneer itself (Eaton et al., 2016; Levy et al., 1992; Truffaut et al, 2017; Pet et al., 2004).

Hybridization between *Q. robur* and *Q. petraea* is quite frequent and adaptive introgression may have played a major role in the latter’s postglacial migration and expansion (Bacilieri et al., 1996; Beatty et al., 2016; Fyfe & Bailey, 1951; Leroy et al., 2017; Leroy et al., 2019; Leroy et al., 2019b; Petit et al., 2004). Despite millennia of hybridization and introgression between these species as well as other European oaks, they maintain morphological and ecological differences (Beatty et al., 2016; Curtu, Gailing & Finkeldey, 2007; Jones, 1959; Kleinschmit, 1993; Leroy et al., 2017; Leroy et al., 2019; Lesur et al., 2019). Controlled pollination experiments suggest that pre- and postzygotic barriers to gene flow in European white oaks maintain the nuclear genetic diversity between species and these barriers are often asymmetric (Lepais et al., 2013; Leroy et al., 2017; Truffaut et al., 2017). Pollen of *Q. petraea* pollinates *Q. robur* more frequently than vice-versa, under both natural and experimental conditions (Lepais et al., 2013; Truffaut et al., 2017). Genomic signatures of admixture suggest that introgression occurs predominantly from *Q. robur* to *Q. petraea* (Guichoux et al., 2012; Leroy et al., 2019b; Petit et al., 2004).

Past studies of genetic differentiation based on small numbers of loci (RAPDs, SCARs, Isozymes, SSRs and AFLPs) reported extremely low differentiation and high genetic homogeneity between *Q. robur* and *Q. petraea* (Barreneche et al., 1996; Bodenes et al., 1997; Coart et al., 2002; Cottrell et al., 2003; Mariette et al., 2002; Scotti-Saintagne et al., 2004; Zanetto et al., 1994), while more recent efforts, based on genome-wide SNP markers (Guichoux et al., 2012; Lang et al., 2018; Leroy et al., 2017; Leroy et al; 2019; Leroy et al., 2019b; Lesur et al., 2018; Reutimann, Gugerli & Rellstab, 2020), have successfully identified some regions significantly differentiated between these two closely related oak species, even though the majority of their genomes still appears permeable to interspecific gene flow.

A common and culturally important habitat for oaks in Britain is parkland. Parklands are wood pastures, often containing deer, that vary in age from 100 to 1000 years (Rackham, 1990). Trees within them can be planted, naturally regenerated, or retained from previous landscapes (Rackham, 1990). Previous studies on the genetic structure of oak populations throughout Britain concentrated on ancient woodland sites, to try to minimise anthropogenic influences and maximise the chances of sampling locally native trees (Cottrell et al., 2002; Petit et al., 2002; Lowe et al., 2004). Chloroplast DNA analyses from ancient woodlands were used to test whether British oak seed stands contain native and locally derived oak (Cottrell et al., 2004; Lowe et al., 2004), and confirmed that most of them were.

In this study we sequenced the whole genome of 386 oak trees from four British parkland sites, including *Q. robur*, *Q. petraea* and their hybrids. We detected over 50 million SNPs and made use of this huge genetic variation to: characterize the structure and evolutionary history of *Q. robur* and *Q. petraea* parkland populations, find loci differentiated between the species, and to detect signatures of recent positive selection in *Q. robur*. We also assembled whole chloroplast genomes and assigned these to haplotypes identified by restriction enzyme fragment sizes in previous studies of oaks in ancient woodlands across Europe, enabling us to test the origins of these parkland oak trees.

## Material and Methods

### Sampling and DNA extraction

Leaf material was collected by the Forest Research Technical Service Unit from 386 oak trees in autumn 2017. Collections were from four British parkland sites: 82 trees in Attingham Park (Atcham, Shropshire, England, 52.688965° N, −2.667944° W), 80 in Hatchlands Park (East Clandon, Surrey, England, 51.257072° N, −0.472332° W), 124 in Langdale Wood (Malvern, Worcestershire, England, 52.085446° N, −2.307670° W) and 100 in Sheen Wood (Richmond, London, England, 51.456229° N, −0.269298° W). Based on morphology, the collectors tentatively identified 376 the trees as *Q. robur* and 10 as *Q. petraea*. Whole genomic DNA was extracted from leaf tissue at RBG Kew using Qiagen DNeasy protocol and was sent to Novogene (Hong Kong) for library preparation and whole-genome shotgun sequencing.

### Genotyping

Shotgun libraries with fragment sizes of 350 bp were prepared with NEBNext DNA Library Prep Kit and were sequenced with 150 bp paired-end Illumina NovaSeq 6000 technology at 22x depth of coverage by Novogene. The quality of the Illumina data generated was assessed with the software FastQC v0.11.5 (Andrews, 2010). Trimmomatic v0.36 (Bolger, Lohse and Usadel, 2014) was used to remove sequencing adapters and for trimming, scanning all reads in windows of four bases, and cutting at the leftmost position if the average Phred base quality score dropped below 20 (99% base accuracy). Reads shorter than 70 bases after trimming were discarded. Processed reads were mapped to the haploid chromosome-level version of the *Q. robur* reference genome (Plomion et al., 2018) with BWA-MEM v0.7.15 (Li & Durbin, 2009) run with default settings. After alignment to reference, PCR duplicates were removed using Samtools v1.9 (Li et al., 2009). Variant calling was performed on all samples simultaneously with the software Haplotype Caller in joint genotyping mode, available in GATK v4.0.8.1 (DePristo et al., 2011; Poplin et al, 2017). SNPs were extracted and filtered to exclude loci with either: quality by depth less than two, Fisher strand test greater than 60, root mean square mapping quality less than 50, mapping quality rank sum test less than −2 or read position rank sum test less than −2. We refer to the resulting SNPs set as the genome-wide set. Summary statistics were computed and plotted using bcftools v1.8.

### Assignment of individuals to species

The genome-wide SNPs set was quality filtered with vcftools v0.1.16 (Danecek et al., 2011) removing loci with missing genotypes and with either: individual mean depth less than 15, minor allele count less than three, minor allele frequency less than 0.005, individual depth less than five or mapping quality less than 20. This set was further reduced by excluding SNPs located in the transposable elements identified by Plomion et al. (2018), with bedtools v2.28.0 (Quinlan & Hall, 2010), and multiallelic sites, with bcftools v1.8. Finally, SNPs were pruned by linkage disequilibrium (r2 > 0.4) using the indep-pairphase function of PLINK v2.0 (Chang et al., 2015) with window size of 50 markers and step of 5. We refer to the resulting SNPs set as the reduced set.

Principal component analysis (PCA) was performed on the reduced set using Plink v2.0 (Chang et al., 2015). We used fastSTRUCTUREv.1.0 (Raj, Stephens, & Pritchard, 2014) to infer admixture levels and to assign individuals to species. fastSTRUCTURE applies a variational Bayesian framework to approximate the model of the widely used STRUCTURE program (Pritchard, Stephens, & Donnelly, 2000). fastSTRUCTURE increases the speed of ancestry inference, while achieving accuracies similar to that of STRUCTURE, by using optimization theory to accelerate the lengthy computation of posterior distributions (Raj et al, 2014). fastSTRUCTURE was run with a simple prior and number of ancestral populations (K) from one to ten. To select the model that best explain structure in the data we used the python utility “chooseK”, available in fastSTRUCTURE, which computes two values for K: one that maximises the marginal likelihood and identifies strong structure and another derived from a heuristic method which is able to identify additional weak structure contributing to variation in the data. Individual trees were assigned to three categories according to the value of the admixture coefficient (q) obtained with fastSTRUCTURE: pure *Q. robur* (q ≥ 0.9), hybrid trees (q ≥ 0.1 & q ≤ 0.9) or pure *Q. petraea* (q ≤ 0.1), as described in Truffaut et al. (2017). We were able to match fastSTRUCTURE designations with species populations on the grounds of an earlier classification of the trees based on leaf morphology. Our collectors’ morphological classification of the trees forcibly assigned sampled individual to either *Q. robur* or *Q. petraea*, as it is difficult to identify hybrids by observation alone (Curtu, Gailing & Finkeldey, 2007; Petit et al., 2004).

### Interspecific differentiation

Genotypes of 10 *Q. petraea* and 10 unrelated *Q. robur* individuals from Attingham park (Atcham, Shropshire, England, 52.688965° N, −2.667944° W) were extracted from the reduced SNP set and were filtered to exclude loci with minor allele frequencies (MAF) below 0.05 using bcftools v1.8. Weir and Cockherham F_st_ (Weir & Cockerham, 1984) was calculated between species using vcftools. We produced F_st_ distribution plots in R (R core team, 2018). We identified outlier loci (1% of most extreme F_st_ values) by generating smoothed quantiles from the empirical distribution of the F_st_-heterozygosity relationship using the R package “fsthet” (Flanagan & Jones, 2017). This method is similar to that described by Beaumont & Nichols (1996) however it avoids assumptions about the underlying population structure and F_st_/Heterozygosity distribution. This is advantageous as population structure parameters are not always known *a priori* and the F_st_-heterozygosity relationship does not always fit the expected distribution and confidence intervals generated with the null infinite island model, particularly when the number of populations is small (< 10), and migration rate is low (Flanagan & Jones, 2017). The method implemented in the package “fsthet” is not a statistical test and always identifies a specific number of outliers from the empirical distribution, depending on the number of loci tested and the chosen confidence interval (Flanagan & Jones, 2017). To mitigate this limitation and minimize the number of false positives we conservatively selected outlier loci showing strong differentiation, with an F_st_ at least two standard deviations from the whole genome mean F_st_. We then used a nonoverlapping 10 kb window approach to estimate the number of outliers per window across the genome, similarly to the method employed in Leroy et al. (2019). We selected the top 1% outlier enriched windows and identified the genes within or flanking (within 5 kb) these regions with bedtools (Quinlan & Hall, 2010), based on the gene models reported by Plomion et al. (2018). To test the species discriminatory power of the identified genes we ran a second fastSTRUCTURE (Raj et al, 2014) taxonomic assignment of all individuals based only on SNP loci located within these genic regions.

### *Population structure and positive selection in* Quercus robur

Genotypes of 360 *Q. robur* individuals identified with fastSTRUCTURE were extracted from the genome-wide SNPs set and were quality filtered with vcftools v0.1.16 (Danecek et al., 2011) removing loci with missing genotypes and with either: individual mean depth less than 15, minor allele count less than three, individual depth less than five or mapping quality less than 20. This set was further reduced by excluding SNPs located in the transposable elements identified by Plomion et al. (2018), with bedtools v2.28.0 (Quinlan & Hall, 2010), and multiallelic sites, with bcftools v1.8. We performed Hardy-Weinberg equilibrium exact test with the function hardy, available in Plink v2.0 and applied the Benjamin and Hochberg correction (Benjamini and Hochberg, 1995) to the resulting p-values, using the R function p.adjust (R Core Team, 2018). We excluded SNPs with p-value less than 0.05 after correction, representing possible genotyping errors. We refer to the resulting SNPs set as the *Q. robur* set.

We used the *Q. robur* set (MAF > 0.05) to estimate linkage disequilibrium decay along the oak genome using two methods: the r^2^ function available in Plink v 2.0 and the tool PopLDdecay (Zhang, Dong, Xu, He, & Yang, 2018). We calculated *Q. robur* nucleotide diversity π (Nei & Li, 1979) along the genome including repeat regions and restricted to genic and intergenic regions, using the vcftools function window-pi in non-overlapping windows of 5 kb.

To study *Q. robur* population structure the *Q. robur* SNP set was pruned by linkage disequilibrium (r2 > 0.4) using the indep-pairphase function of PLINK v2.0 (Chang et al., 2015) with window size of 50 markers and step of 5. PCA (Plink v2.0) and fastStructure with number of ancestral clusters K from one to ten were performed on the pruned *Q. robur* set excluding loci with minor allele frequencies below 0.05.

Marker based realized genomic relatedness was computed across sites including all sampled individuals and restricted to each site, according to the formula by VanRaden (2008), implemented in the kin function of the R package synbreed (Wimmer, Albrecht, Auinger, & Schon, 2012), using the pruned *Q. robur* SNPs set (MAF > 0.05). Related individuals were excluded by removing an individual from each pairwise comparison with realized relatedness (VanRaden, 2008) greater than 0.05. PCA was re-computed in Plink v2.0 for the unrelated *Q. robur* individuals. We identified potential parent-offspring and siblings relationship based on our relatedness estimates and the approximate age of trees derived from diameter at breast height.

We used SweeD v3.2.1 (Pavlidis, Živković, Stamatakis, & Alachiotis, 2013) and OmegaPlus (Alachiotis, Stamatakis, & Pavlidis, 2012) to perform selective sweep scans of the *Q. robur* genome, including repeat regions and all allele frequencies but excluding multiallelic loci and loci with missing calls. Hard sweeps are drastic forms of selection characterized by the appearance of a new beneficial mutation with a strong selective advantage in a population which quickly increases in frequency until it reaches total fixation (Nielsen, 2005; Weigand and Leese, 2018). This process generates a distinct signature in the genomic region surrounding the selected variant: a decreased overall nucleotide diversity due to the genetic “hitchhiking” effect, an increase in low and high frequencies variants and distinct linkage disequilibrium patterns (Nielsen, 2005; Weigand and Leese, 2018). Following fixation of the beneficial allele in a population, mutation and recombination re-increase diversity in the surrounding region and slowly degrade the patterns typical of hard sweeps (Nielsen, 2005; Weigand and Leese, 2018). SweeD employs a composite likelihood test to detect hard sweeps based on the site-frequency spectrum (SFS) of SNPs in whole-genome data.

OmegaPlus searches for specific linkage-disequilibrium patterns characteristics of recent selective sweeps and output the ω-statistic. SweeD and OmegaPlus were run with a grid parameter that resulted in a measurement of CLR and ω-statistic every 5,000 bp and each chromosome was scanned separately. In OmegaPlus the minimum and maximum size of the sub-region around a position which was included in the calculation of the ω-statistic were fixed to 500 and 100,000 base pairs, respectively. Ancestral allele states for the SweeD analysis were inferred using two outgroups, *Fagus sylvatica L.* (Mishra et al., 2018) and *Castanea mollissima* (Xing et al., 2019). Raw whole genomic sequencing reads for both outgroups were downloaded from the European Nucleotide Archive (ENA) (PPRJEB24056, RJNA527178) and were mapped to the *Q. robur* reference genome using Bowtie2 in local alignment mode. The alignments generated were sorted and processed with Samtools v1.9 (Li et al., 2009) to remove PCR duplicates. Outgroups genotypes, including homozygous reference calls, were inferred with bcftools v1.8 and the vcftools v0.1.16 (Danecek et al., 2011) utility “fill-aa” was used to record the ancestral state for the *Q. robur* SNPs. The ancestral allele was determined only for SNP loci which appeared homozygous for the same allele in both outgroups. If a site was either not covered by the outgroup reads, was heterozygous in one or both outgroups or the outgroups were homozygous for alleles not found in oaks, then the ancestral state for that locus was not determined. We identified the common outliers, 1% of most extreme p-values, between the SweeD and OmegaPlus runs. We used bedtools to identify the gene models (Plomion et al., 2018) within or flanking the common outlier windows.

### Population size history and species divergence

We inferred *Q. robur* and *Q. petraea* population size and separation history using a multiple sequentially Markovian coalescent approach implemented in the software MSMC (Schiffels & Durbin, 2014). MSMC is an extension of the pairwise sequential Markovian coalescent (PSMC) model (Li & Durbin, 2011), which analyses two homologous sequences from a diploid individual, therefore limiting the resolution of population size estimation of more recent history (Schiffels & Durbin, 2014). MSMC employs a simplification which extends PSMC to multiple sequences (Schiffels & Durbin, 2014). For population size history inferences, we ran MSMC on one, two and four high coverage (mean depth > 20x) individuals of each species separately, to achieve higher resolution for both distant and recent history, respectively. For species separation history we used four individuals, two *Q. robur* and two *Q. petraea* trees. Biallelic genotypes were extracted from the genome-wide set, excluding loci with missing data, and MSMC was run over the 12 chromosomes with default time segment patterning. Mappability mask files of repeat elements (Plomion et al., 2018) were provided in bed format to exclude non-unique regions of the genome from the analysis. Statistical phasing was performed using Beagle 5.1 with default settings (Browning & Browning, 2007). MSMC outputs times and population sizes scaled by mutation rate per base pair per generation. To convert the output to real time and sizes it is necessary to divide estimates by the mutation rate and further multiply these by the generation time. As the mutation rate of *Q. robur* and *Q. petraea* was unknown, we used the substitution rate 7.5 × 10^−9^ of *A. thaliana,* retrieved from Buschiazzo, Ritland, Bohlmann, & Ritland (2012), as it was done for *Fraxinus excelsior* in Sollars et al. (2017). We used a generation time of 50 years (Leroy et al., 2019; Leroy et al., 2020). Effective population size history and separation time estimates were plotted in R (R Core Team, 2018).

### Chloroplast Phylogeny

The chloroplast DNA sequences of 309 of the 386 individual trees were assembled *de novo* using Novoplasty 3.7.2 (Dierckxsens, Mardulyn & Smits, 2017). The software was run in chloroplast mode with k-mer length of 39, the minimum length of overlap necessary to join adjacent reads in the assembly. We used the *Zea mays* chloroplast gene for the large subunit of ribulose bisphosphate carboxylase (RUBP) as seed sequence for the assembly and the *Quercus lobata* isolate SW786 complete chloroplast sequence (Sork et al., 2016) as reference. The assembly was performed *de novo* by Novoplasty with the reference used to aid resolution of difficult regions, such as inverted repeats. Novoplasty was set with read length of 150 and insert-size of 350 and the software was allowed to automatically finetune these values.

Haplotype networks are often used to visualize intraspecific phylogenies which can be challenging to reconstruct due to low variation between individuals; this is accentuated in non-recombining sequences such as chloroplast DNA. The median joining method has been proposed as alternative to maximum likelihood methods to build networks that best express the large number of plausible trees in such cases and is able to handle large dataset efficiently, as it is often the case with intraspecific data (Bandelt et al., 1999). The median joining method described by Bandelt et al. (1999) can be perceived as an intermediate method between the minimum spanning tree algorithm by Kruskal (Kruskal, 1956) and Farris’s maximum parsimony algorithm (Farris, 1970) and it is applicable to nucleotide sequences.

The 309 fully assembled and circularized chloroplast sequences were linearized and shifted to the same origin, which was set at the seed RUBP sequence, using the Perl package fasta-tools (https://github.com/b-brankovics/fasta_tools). The shifted complete chloroplast sequences were aligned using MAFFT v7 (Katoh, Misawa, Kuma, & Miyata, 2002; Katoh & Standley, 2013) and the alignment was exported in Phylip format. PopArt v1.7 (Leigh & Bryant, 2015) was used to generate a median joining network from the alignment file as described by Bandelt, Forster, & Rohl (1999).

### Chloroplast haplotypes identification

Representative chloroplast DNA sequences for each of the clusters identified in the median joining network were analysed by reproducing *in silico* the polymerase chain reaction-restriction fragment length polymorphism (PCR-RFLP) method used to characterize the main oak chloroplasts lineages in Europe, including over 40 haplotypes, in Petit et al. (2002b). The haplotypes identified in our chloroplast network were matched to known oak chloroplasts haplotypes according to the information provided by four non-coding chloroplast DNA fragments extracted and digested *in silico* with custom scripts. The four chloroplast DNA fragments and restriction enzyme pairs were: trnD-trnT (DT) with TaqI, psaA-trnS (AS) with HinfI, psbC-trnD (CD) with TaqI and trnT-trnF (TF) with AluI (Petit, Demesure, & Dumolin, 1998; Petit et al., 2002b). The first three primers/restriction enzyme pairs were first described in Demesure, Sodzi and Petit (1995) while the latter was first described in Taberlet, Gielly, Pautou and Bouvet (1991). The length variants obtained with restriction digestion simulation of the chloroplast DNA fragments were compared and matched with those reported for previously characterized oak haplotypes (Appendix B in Petit et al., 2002b). The trnD-trnT and trnT-trnF fragments were further characterized for the presence of a point mutation, using AluI and CfoI restriction enzymes respectively, as described in Petit et al. (2002b).

We compared the distribution of the chloroplast haplotypes identified with that of 178 ancient woodlands across England, surveyed in a previous study (Cottrell et al., 2002). To assess whether the parklands surveyed matched the local ancient woodlands dominant haplotype, we performed an ordinary kriging linear regression (R, “gstat” package) of Cottrell et al. (2002) data to define haplotypes dominance regions in Britain, similarly to Lowe et al. (2004).

## Results

### SNP discovery

We sequenced the whole genome of 386 oak trees from British parklands and identified 59,310,788 SNPs by mapping to the *Q. robur* reference genome (Plomion et al., 2018). Mean individual depth of coverage of SNP loci was 21.5x, varying from 12.7x to 33.4x across samples. We identified an average of 8,083,170 sites differing from the reference genome per sampled tree (Plomion et al., 2018) and missing genotypes varied from 1 to 5% per individual, with mean of 4.2%. The Ts/Tv ratio was normally distributed around mean 2.58 with small variation, as expected. In total 4,838,132 SNPs (~8%) were in genic sequences, according to the gene models reported by Plomion et al. (2018).

### Population structure and interspecific differentiation

We analysed the genome-wide variation of 386 oak trees with principal component analysis (PCA) and model-based inference based on the reduced SNPs set, which included 2,768,547 unlinked SNPs (r^2^ < 0.4). The first principal component explained 38% of the total variance and clearly separated *Q. robur* and *Q. petraea* individuals (Figure 1). The second principal component accounted for 10% of the variance and separated the two species to a lesser extent. The explained variance levelled off at the third component (Figure 1C).

**Figure 1.**
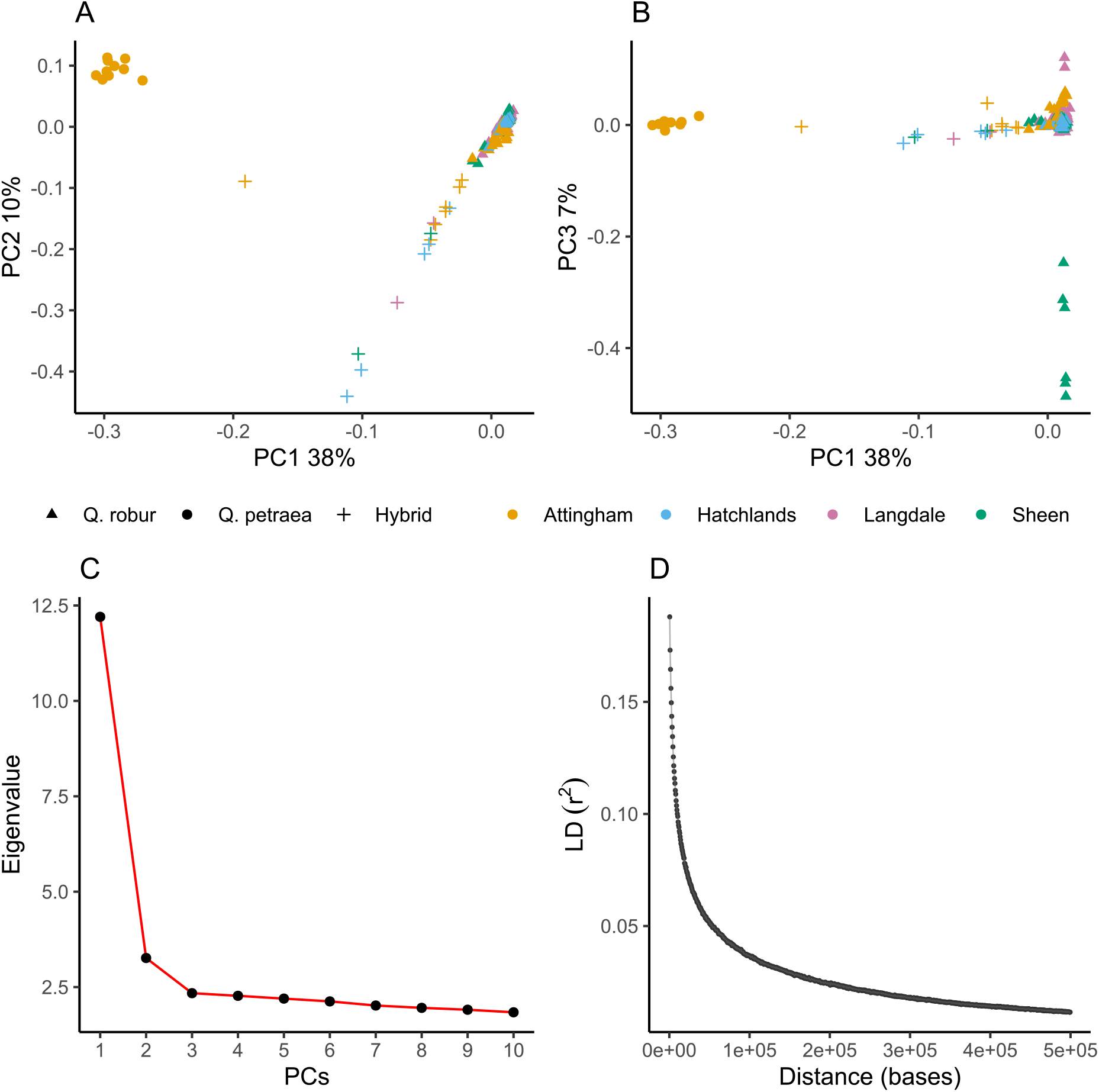
PCA of 386 individual oak trees of two species, based on 2,768,547 unlinked SNPs (r^2^ < 0.4). A) PC1 against PC2. B) PC1 against PC3. C) Eigenvalues of the computed principal components. D) LD decay in the *Q. robur* genome: points are 500 bases apart.

Our fastSTRUCTURE Bayesian models with two different metrics concordantly selected K=2 as the number of ancestral clusters that maximises the marginal likelihood of our data (Raj et al., 2014). In total, 10 individuals were assigned to *Q. petraea*, 360 individuals to *Q. robur* and the remaining 16 individuals were classified as hybrids (Figure 2). The fastSTRUCTURE species assignment matched the collectors’ classification based on leaf morphology for pure individuals, except for three *Q. petraea* individuals previously assigned to *Q. robur*. In a PCA of all individuals the admixed individuals came between the main species clusters (Figure 1A). Three of the 16 admixed individuals identified were originally classified morphologically as *Q. petraea*, while the remaining 13 had been assigned to *Q. robur.* The collectors’ classification of the identified admixed individuals reflected the dominant contribution to their ancestry (Figure 2).

**Figure 2.**
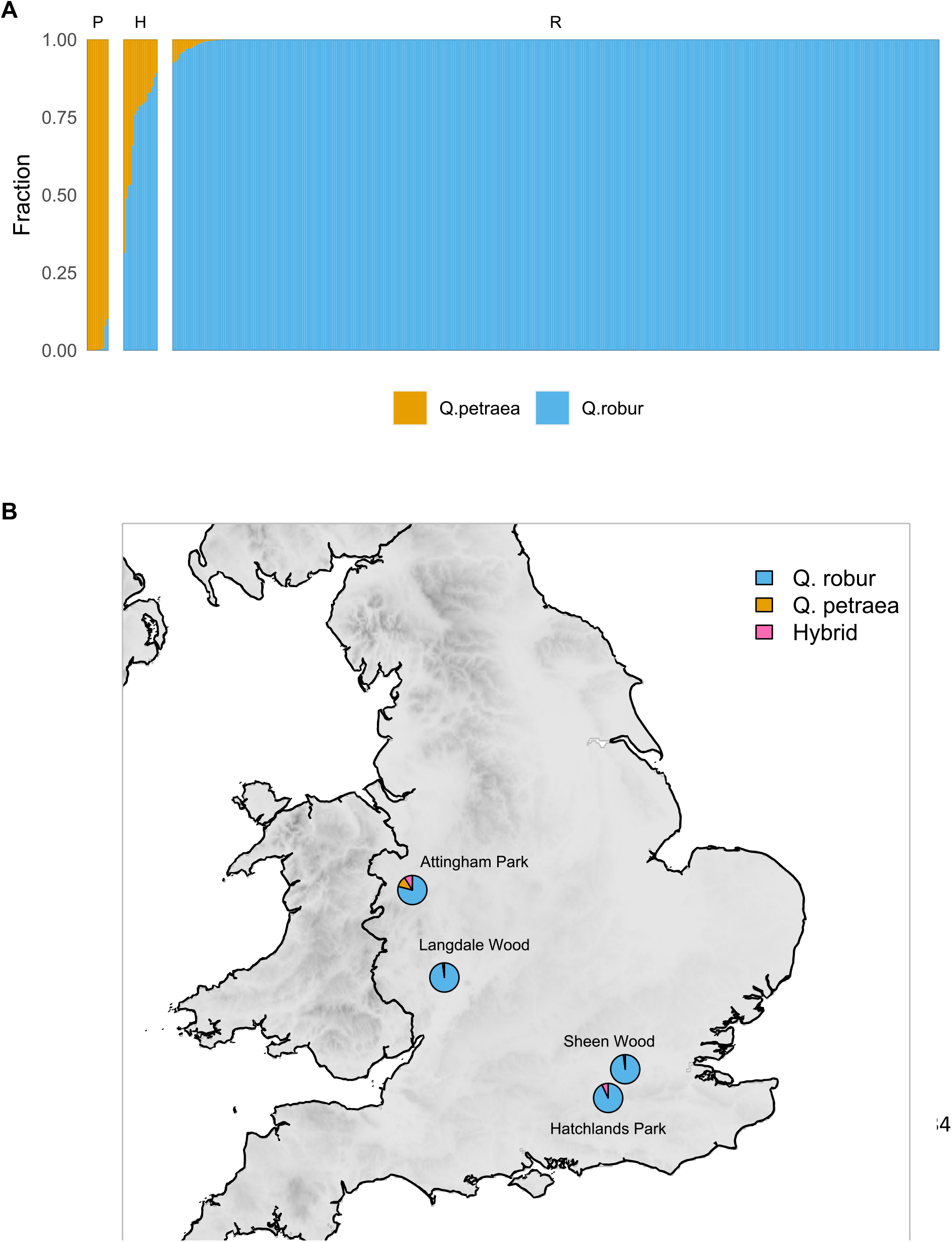
A) Bar plot of fastSTRUCTURE results of 386 individuals based on 2,768,547 unlinked SNPs (r^2^ < 0.4), K = 2. Q = *Q. robur*, P = *Q. petraea*, H = Hybrid. B) Map of species distribution across sampling sites.

Between ten unrelated *Q. robur* and the ten *Q. petraea* individual trees from Attingham park loci with minor allele frequencies over 0.05 (914,242 SNPs, ~24% genic and ~76% intergenic) gave a mean interspecific F_st_ of 0.158 (median and standard deviation were 0.111 and 0.156, respectively). Those SNPs located in the gene models predicted by Plomion et al. (2018) (223,319 SNPs), gave a mean F_st_ value of 0.155 (median and standard deviation were 0.111 and 0.153, respectively) whereas intergenic SNPs gave a mean F_st_ value of 0.159 (median and standard deviation were 0.111 and 0.157, respectively). The genome-wide F_st_ L-shaped distribution shows that the majority of loci exhibit small differentiation (Figure S2, Supporting Information). A minority of loci clearly displayed very strong differentiation and 610 SNPs were fixed for different alleles in the two species (F_st_ = 1). Within-species F_st_ calculation between two populations (Attingham and Hatchlands) each composed of 10 unrelated *Q. robur* individuals gave a much lower mean F_st_ value of 0.024 (median and standard deviation were 0 and 0.044, respectively).

Between *Q. robur* and *Q. petraea* we identified 8019 outlier SNPs, representing most extreme values of the F_st_-heterozygosity distribution (Figure S3, Supporting Information). We searched the *Q. robur* gene models (Plomion et al., 2018) within or flanking the top 1% of outlier-enriched 10 kb windows (69 windows with 5 or more outliers) and found 81 genes (Table S1, Supporting Information). Using only these 81 genic regions in fastSTRUCTURE provided the same assignment of individuals to species as the runs based on 2,768,547 genome-wide SNPs, except that three individuals previously classified as *Q. robur* with signals of introgression from *Q. petraea* were classified as hybrids with a strong Q. robur component.

### Quercus robur *genetic diversity and positive selection*

The PCA restricted to the *Q. robur* individuals, based on 839,911 unlinked SNPs (MAF > 0.05) showed that there is little difference in the variance explained by each principal component and there is no major cluster visible in the plots (Figure S4, Supporting Information). Realized genomic relatedness (VanRaden, 2008) among the 360 *Q. robur* trees was normally distributed around 0 (Figure S5, Supporting Information), as expected, and relatively low with 123 pairwise estimates above 0.125 (expected for third-degree relatives) among 64,620 pairwise estimates. There were significant differences in relatedness within sites (Figure S6, Supporting Information). Trees within Hatchlands Park had the lowest level of relatedness (mean = 0.0016) with only 7 pairwise estimates above 0.125, involving 14 trees out of the 75 *Q. robur* sampled at this site (18 %). Sheen Wood similarly had relatively low relatedness levels (mean = 0.0019) with 23 estimates above 0.125, involving 21 unique trees out of 98 (21%). In contrast, Langdale Wood (mean = 0.003) and particularly Attingham Park (mean = 0.0065) reported significantly higher relatedness levels, with 48 and 45 pairwise estimates above 0.125 respectively, involving 53 unique trees (43%) at Langdale Wood and 40 unique trees (61%) at Attingham Park. Based on our relatedness estimates and the approximate age of trees based on diameter at breast height, we identified two potential parent-offspring relationships and two full-sibs pairs at Attingham Park, three full-sibs pairs at Sheen Wood, five full-sibs pairs at Langdale Wood and a single potential full-sib relationship at Hatchlands Park. In a global test of relatedness among sites, we found only two among-site pairs with relatedness above 0.05. These were between Hatchlands Park and Sheen Wood.

Examining genomic variability of *Q. robur* among all four populations with PCA, after excluding 99 individuals to remove any relatedness relationship above 0.05 between samples, PCA eigenvalues levelled off reflecting the lack of any strong geographically correlated structure in the nuclear genomes of *Q. robur* individuals between the four parklands sampled (Figure S7, Supporting Information). Running fastStructure on the 360 *Q. robur* individuals (K from one to ten) also confirmed this lack of structure within *Q. robur*: both metrics employed for the choice of ideal model complexity concordantly suggested a single ancestral population.

Linkage disequilibrium (LD) was found to decay quickly in the 360 *Q. robur* individuals, with average r^2^ below 0.2 at 500 bases apart (Figure1, Figure S8, Supporting Information). Over 90% of LD-blocks size estimates are within 5,000 bp length (Figure S8, Supporting Information). In windows of 5,000 bp, *Q. robur* pairwise nucleotide diversity (Figure 3) showed genome-wide diversity π of ~0.007, with little variation between chromosomes and parklands sites. Nucleotide diversity was higher in the intergenic regions (π = 0.0066) than the coding regions (π = 0.0024), as expected.

**Figure 3.**
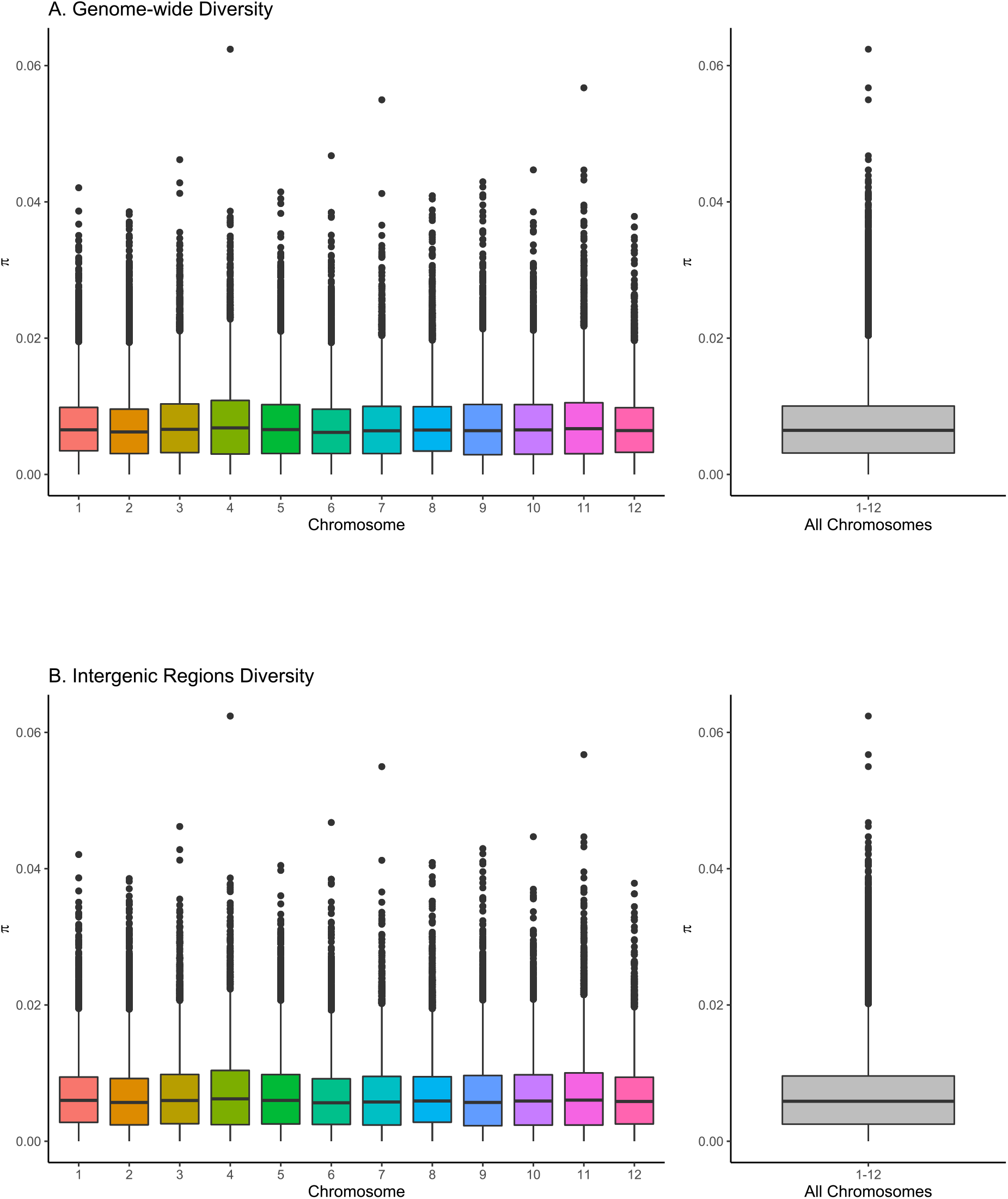
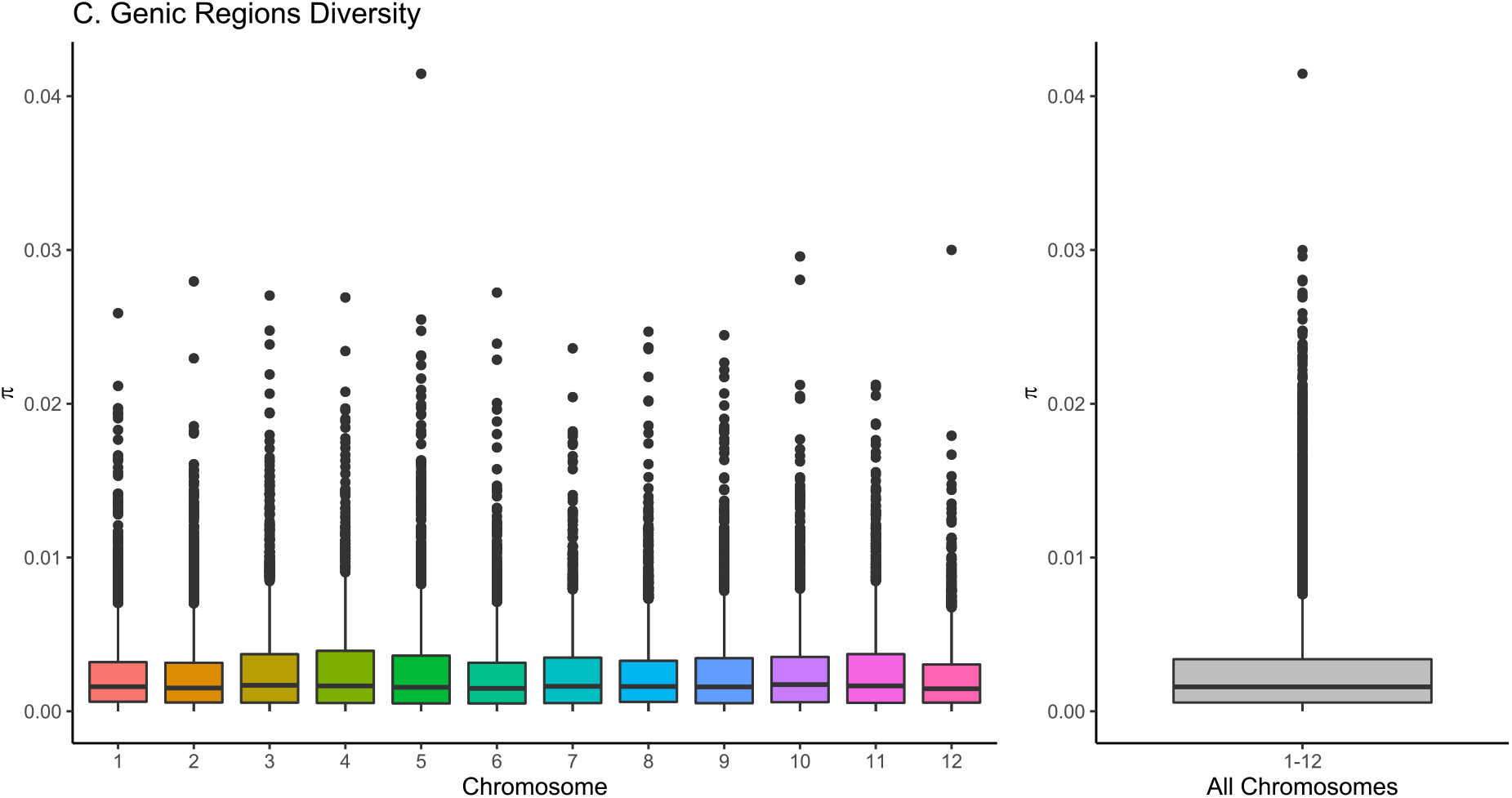
Pairwise nucleotide diversity π bar plots computed in windows of 5,000 bp across the Q. robur genome, based on 360 individuals and 59,310,788 SNPs.

Selective sweep scans every 5,000 bp along the 12 *Q. robur* chromosomes using SweeD (Pavlidis et al, 2013) and OmegaPlus (Alachiotis et al., 2012) each identified 1,311 outlier regions (1% of most extreme values). There were 10 common outlier regions found by both methods (p < .01) (Figure S9, Supporting Information), located on chromosomes 1 (2 regions), 2, 3 (2 regions), 9, 11 (3 regions) and 12. The gene models (Plomion et al., 2018) within or flanking the identified sweep regions (Table S2, Supporting Information) included key developmental and stress response regulators, such as: hydrophobic seed proteins, which are expressed on the seed surface and appear to play a role in seed survival by regulating water-uptake and affecting pathogen attachment and penetration (Gijzen et al., 1999); zinc-finger proteins, transcription factors that regulate plant growth and are thought to be involved in the response to environmental stresses such as salinity, cold and draught (Han et al., 2020; Huang, Wang, & Zhang, 2004); tyrosine kinases, key transmembrane receptors involved in signal transduction and linked to plant growth and response to both biotic and abiotic stresses (Miyamoto et al., 2019); copines, a class of calcium-dependant phospholipid-binding proteins linked to disease resistance and acclimation in *A. thaliana* and *Triticum aestivum* (wheat) (Jambunathan, Siani, & McNellis, 2001; Jambunathan & McNellis, 2003; Zou et al., 2016; Zou, Ding, Liu, & Hua, 2017); Vacuolar H+-ATPase, a proton pump found in organelle membrane involved in ion, metabolites and pH homeostasis in plants (Dettmer, Hong-Hermesdorf, Stierhof, & Schumacher, 2006) that appears to cover a crucial role in plant tolerance to salt induced stress (Golldack & Dietz, 2001; Padmanaban et al., 2004; Ratajczak, 2000; Zhang et al., 2012); patellin proteins, phosphatidylinositol membrane transfer proteins seemingly involved in development, response against salt stress and immunity against certain viruses in plants (Peiro et al., 2014; Zhou et al., 2019).

### Species population size and separation history

Multiple sequentially Markovian coalescent (MSMC) plots for both species suggest a long-term history of population size decline followed by a steep increase in more recent history (Figure 4). Using 2, 4 or 8 nuclear haplotypes gave similar results for time periods where their estimates overlap (Figure 4A-C).

**Figure 4.**
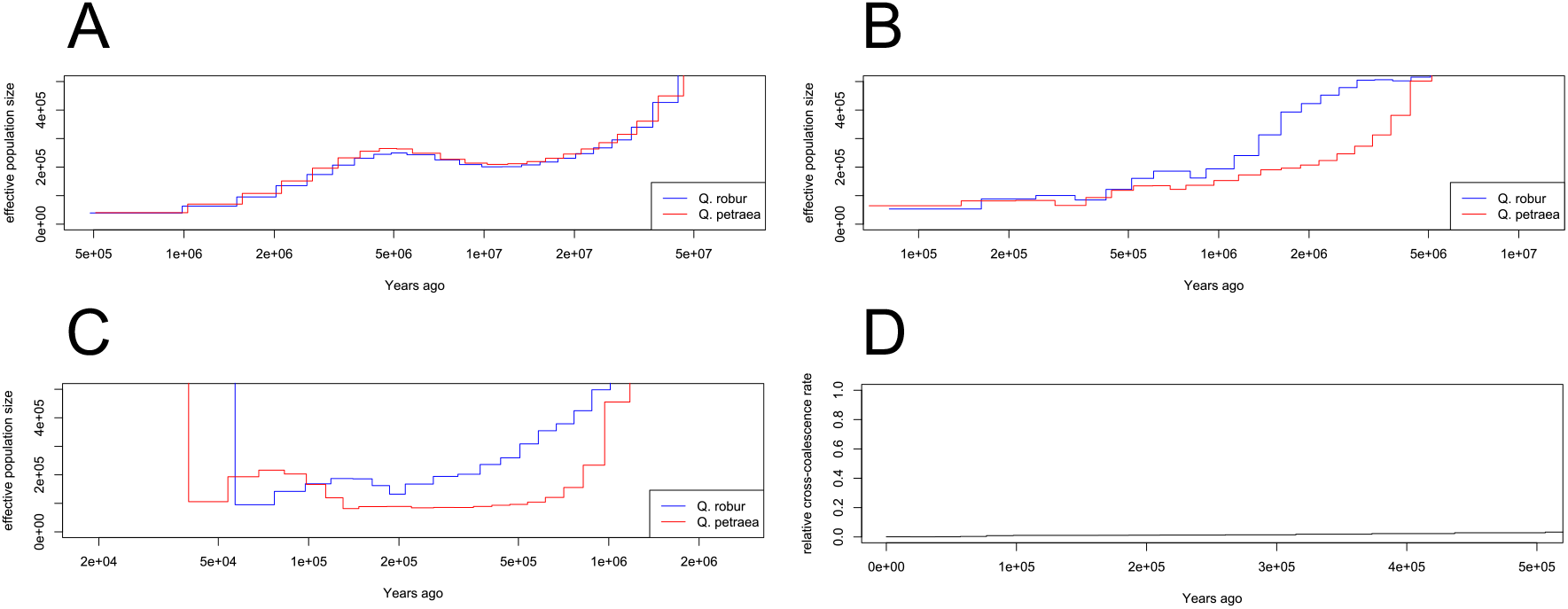
Effective population size and separation history of *Q. robur* and *Q. petraea* estimated with the MSMC method. Generation time: 50 years. Mutation-rate:7.5e^−9^. A) Estimates based on 2 haplotypes for each species. B) Estimates based on 4 haplotypes for each species. C) Estimates based on 8 haplotypes for each species. D) Relative cross-coalescence rate (CCR) ratio per year. The CCR ratio is a measure of divergence and represents the ratio of between-populations over within-populations coalescence rate. Values close to 0 indicate that populations have diverged, while values close to 1 indicate that populations have not yet diverged as between and within populations coalescence rates are equal.

Estimation of cross-coalescence rates with MSMC suggests that *Q. robur* and *Q. petraea* populations were already separated 500 Kya (Figure 4D).

### Chloroplast haplotypes phylogeny and identification

We *de novo* assembled and circularized the full chloroplast DNA sequence of 309 oak samples, including 287 *Q. robur*, 8 *Q. petraea* and 14 hybrids. A median-joining plastid haplotype network identified five major chloroplast haplotypes, which we arbitrarily labelled from I to V, with lengths varying from 161,148 bp to 161,306 bp (Figure 5). We matched the five haplotypes to those previously identified using the PCR-RFLP method (Petit et al. 2002b). Haplotypes I to IV displayed a particular point mutation in the trnD-trnT chloroplast fragment (Table S3E) characteristic of haplotypes belonging to lineage “B”, which hypothetically originated in a late Pleistocene glacial refugium located in the Atlantic side of the Iberian Peninsula (Cottrell et al., 2002; Petit et al., 2002a; Petit et al., 2002b). Similarly, Haplotype V exhibited a particular point mutation in the trnT-trnF chloroplast fragment (Table S3E) characteristic of haplotypes stemming from lineage “A”, which originated in a Pleistocene glacial refugium located in the Balkans geographic area (Cottrell et al., 2002; Petit et al., 2002a; Petit et al., 2002b).

**Figure 5.**
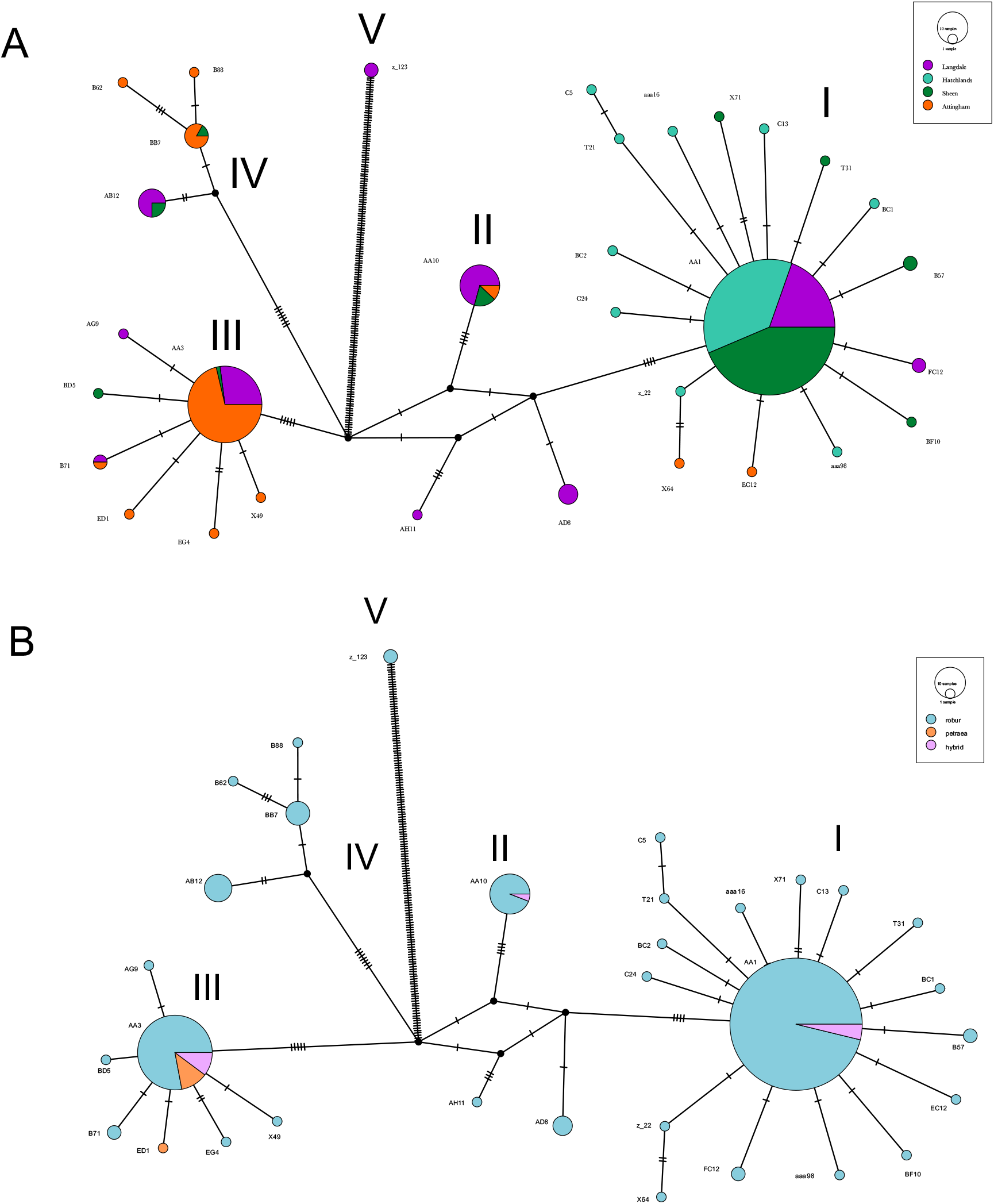
A) Median-joining chloroplast haplotype network of 287 *Q. robur,* 8 *Q. petraea* and 14 hybrid individuals. Colours represent sites and the size of nodes is proportional to number of individuals. Hatch marks along edges represent the number of single base mutations between nodes. Edges length does not carry any weight. The major nodes identified were arbitrarily labelled from I to V. B) Colours represent species.

Haplotypes I and III were matched with haplotypes 10 and 12, respectively (Table S3), described in Petit et al. (2002b), according to the information provided by restriction digestion of the DT chloroplast fragment (Table S3A). Haplotype II in our study displayed a 13 bp duplication in the TF chloroplast DNA fragments (Table S3D) characteristic of haplotype 11 (Cottrell et al., 2002). Haplotypes III and IV represents variations of haplotype 12, labelled as 12a and 12b in previous surveys (Cottrell et al., 2002; Petit et al., 2002a; Petit et al., 2002b). Haplotype V in our study was matched with either haplotype 7 or 26, two closely related Balkans haplotypes, according to the information provided by digestion of the DT and AS fragments (Table S3, Supporting Information).

The two predominant haplotypes identified across the four sampled sites were haplotype 10 and 12 (Figure S10, Supporting Information). Haplotype 10 is predominant at Sheen Wood and Hatchlands Park in the south-east. It is also well-represented at Langdale Wood, however it is almost absent from Attingham Park, which is located further north-west in Shropshire. Differently, haplotype 12 is nearly absent from the south-eastern sites while it is present at Langdale Wood and it is the predominant haplotype at Attingham Park.

According to our ordinary kriging linear regression based on Cottrell et al. (2002) ancient woodlands data (Figure S11, Supporting Information), Attingham Park (dominant haplotype 12) and Langdale Wood (dominant haplotype 10) did not occur in an area dominated (> 50%) by a single haplotype, but both sites are located in close proximity ( < 25 km) to a matching dominant autochthonous haplotype patch. Hatchlands Park (dominant haplotype 10) and Sheen Wood (dominant haplotype 10) do not have the same dominant haplotypes as local ancient woodlands according to our interpolation, however this area of England appears particularly fragmented and dominance regions for all three major British haplotypes (haplotypes 10,11 and 12) are found in proximity (~50 km) to both sites (Figure S11, Supporting Information). There was no distinction between species: most of the *Q. robur* individuals identified at Attingham Park shared the same haplotype of the *Q. petraea* individuals from the same site (Figure S10, Supporting Information).

## Discussion

### Population structure and Interspecific differentiation

We sequenced the whole genome of 386 oak trees across four British parkland sites and characterized over 50 million SNPs in British *Q. robur* and *Q. petraea*. We found evidence for hybridisation between the two species and found this to be slightly more common than suggested by morphology alone. Trees with evidence for hybridisation in their genomes generally exhibited predominantly the morphological traits of one of the parental species and had thus been previously identified as one parental species on the basis of morphology alone. Previous studies have found that oak hybrids often display uneven combinations of phenotypic characters of the parental species, especially where there has been recurrent backcrossing (Curtu, Gailing & Finkeldey, 2007; Petit et al., 2004). Three individuals could represent first-generation (F1) hybrids in our samples, and the others showed evidence of backcrossing to *Q. robur*, likely due to the predominance of this species in the parklands surveyed.

The genomes of *Q. robur* and *Q. petraea* were largely undifferentiated, confirming previous results from sites across Europe (Barreneche et al., 1996; Bodenes et al., 1997; Coart et al., 2002; Lesur et al., 2018; Mariette et al., 2002; Saintagne et al., 2004; Scotti-Saintagne et al., 2004; Zanetto et al., 1994). We reported mean F_st_ of 0.155 between *Q. robur* and *Q. petraea* coding regions, close to recent estimates (mean F_st_ = ~0.13) by Lang et al. (2018), who studied populations in central and western Europe. We did not find any significant difference between intergenic and genic F_st_ trends. We identified 81 genic regions, with strong species discriminatory power. Two of these genic regions were previously found in a study of genome-wide differentiation among four white oak species (*Q. robur*, *Q. petraea*, *Q. pyrenaica* and *Q. pubescens*) in France, which identified a total of 215 regions of differentiation, 133 of which contained genes (Leroy et al., 2019).

### *Nucleotide diversity and positive selection in* Q. robur

Linkage disequilibrium was found to decay rapidly in *Q. robur* and we reported genome-wide nucleotide diversity π of 0.007 in this species and our coding region estimates (π = 0.0024) are not far from those reported in Lang et al. (2018) for *Q. robur* and *Q. petraea*, 0.0037 and 0.0039, respectively, calculated across over 800 genic regions and based on 25 individuals from stands in central and western Europe.

The differing levels of relatedness that we found among trees within parklands could reflect different management practices in different parklands in the past. Trees in Attingham Park had particularly high relatedness and we identified two potential parent/offspring relationships, two pairs of full-sibs and numerous second- and third-degree relatives there. Trees in Langdale Wood and Sheen Wood had slightly lower levels of relatedness however potential full-sibs (five at Langdale Wood and three at Sheen Wood) and several second- and third-degree relatives can be found at each site. Hatchlands Park showed significantly less kinship. It is likely that sites with higher levels of relatedness had high natural regeneration in the past, and perhaps planting of locally sources acorns.

We identified 10 genes with, or in proximity to, signatures of recent positive selection in *Q. robur* based on both site frequency spectrum and linkage disequilibrium patterns. The putative functions of these suggest that they may be involved in stress tolerance. None of them were within the top 81 F_st_ outlier regions strongly differentiated between *Q. robur* and *Q. petraea*. However, one of these regions (Qrob_T0000290.2, hydrophobic seed protein, Table S2 Supporting Information) did display moderate levels of interspecific differentiation (mean F_st_ = 0.47).

### Population history and secondary contacts

Hybridization and extensive exchanges of chloroplast DNA between European oaks have been well documented (Dumolin-Lapegue, Kremer, & Petit, 1999; Lepais et al., 2009; Leroy et al., 2019; Petit et al., 2002b; Petit et al., 1997) and most recent genetic studies based on ABC models have shown that massive secondary contacts between European white oak species have occurred recently after a long period of isolation, probably at the end of the last glacial period or at the start of the current interglacial period (Leroy et al., 2017; Leroy et al., 2019). These contacts are thought to have homogenized the majority of the nuclear genome of European white oak species with the exception of some barrier regions accumulated during the long periods of isolation that preceded secondary contacts (Lepais et al., 2013; Leroy et al., 2019; Petit et al., 2002a). Our estimates of past effective population sizes suggested that *Q. robur* and *Q. petraea* shared a similar history of population size decline during the last 2.5 million years followed by a recent postglacial re-expansion. Due to uncertainties about generation time and mutation rates in oak, we do not know if the onset of this apparent decline was when Eurasian white oaks colonised Europe from America, or if it began due to Pleistocene glaciations (Cottrell et al., 2002b; Leroy et al, 2017; Mazet et al., 20915; Petit et al., 2002a; Petit et al., 2002b). The steep increase in population sizes recorded in more recent history in both species using 8 haplotypes seems to reflect the post-glacial expansion of white oaks in Europe that is reported to have taken place after the last glacial maximum (Cottrell et al., 2002; Leroy et al, 2017; Petit et al., 2002a; Petit et al., 2002b). Our estimation of cross-coalescence rates is concordant with the hypothesis that despite the extensive interspecific gene-flow in the late Pleistocene, the species were already fully diverged 500 Kya (Hipp et al., 2019; Leroy et al., 2017; Leroy et al., 2019).

### British parkland chloroplast haplotypes

We detected four native British oak chloroplast haplotypes (Haplotypes 10, 11, 12 and 7) in the parklands we studied, and no haplotypes that could be attributed to planting of non-native seed stocks. The vast majority of our samples (> 99%) possess haplotypes thought to be derived from Iberian glacial refugia, but two individuals at Langdale Wood displayed haplotype 7 (Lineage “A”), from a Balkan refugium. This Balkan haplotype is rare in Britain, but has had a long presence in the Forest of Dean and surrounding area, where it is the dominant haplotype of the oldest trees group (133-281 years old) and has been detected in a 320-year-old tree (Cottrell et al., 2002; Cottrell et al., 2004; Lowe et al., 2004). Langdale Wood, where we identified this haplotype, is located approximately within 50 km from the Forest of Dean. It is not known if this haplotype reached the Welsh marches naturally from the Balkans through natural postglacial colonization or if human-mediated activity, such as the return of medieval knights from crusades, is responsible for the scattered presence of this haplotype in the north of France and around the Forest of Dean (Cottrell et al., 2002; Cottrell et al., 2004).

The chloroplast haplotypes present in the four parklands fit well with local ancient woodland haplotype distributions (Cottrell et al., 2002; Lowe et al., 2004). According to our kriging interpolation of dominant haplotypes in Britain, based on the Cottrell et al. (2002) data, all four parkland sites surveyed are located within a short radius (25-50 km) of a matching autochthonous dominant haplotype region (Figure S11, Supporting Information). This suggests that these parkland oaks derive from local seed sources, but we cannot fully exclude the possibility that seeds could have been imported from different regions of Britain with the same dominant haplotype, or even from Spain or France where these haplotypes are similarly abundant (Petit et al., 2002; Lowe et al., 2004).

## Acknowledgements

We thank Forest Research and its Technical Service Unit for sending us leaf samples. We thank Joan Cottrell (Forest Research) for sharing with us haplotype data of 178 ancient oak woodlands. We thank Richard Nichols for his expert advice on our statistical analyses. We thank Alberto Carmagnini (Queen Mary University of London, Laurent Frantz’s Lab) for his constructive comments and assistance in various bioinformatics analyses. Finally, we thank all woodland owners, managers and staff that allowed us and our collaborators to access and sample trees at their sites. This research utilised Queen Mary’s Apocrita HPC facility, supported by QMUL Research-IT: http://doi.org/10.5281/zenodo.438045.

This project is part of a PhD programme run jointly between Queen Mary University of London and the Royal Botanic Gardens, Kew funded by the UK Government Department for Environment, Food and Rural Affairs (Defra), in association with Action Oak. Sequencing was funded by Defra under the Future Proofing Plant Health scheme.

## Data Accessibility

DNA sequences: trimmed sequencing reads have been deposited in the European Nucleotide Archive under the Project Accession no. PRJEB30573. Samples metadata available in the supplementary spreadsheet provided.

## Author contributions

Gabriele Nocchi analysed the data and wrote the manuscript, Nathan Brown sampled and phenotyped the trees, Tim Coker extracted DNA, William Plumb extracted DNA, Jonathan Stocks extracted DNA, Sandra Denman selected the sites and organised the sampling, Richard Buggs obtained funding, oversaw the project and helped write the manuscript.

