## Supporting Information for "Genomic structure and diversity of oak populations in British Parklands"

**Table S1.** Summary table of the 81 gene models within or flanking the top 1% $F_{st}$ outlier-enriched 10kb windows between *Q.robur* and *Q.petraea*.

| ID | Type | Start | End | Score | Gene ID |
|---|---|---|---|---|---|
| Qrob_Chr01 | mRNA | 24923692 | 24927355 | 100.0 | ID=Qrob_T0001470.2 |
| Qrob_Chr01 | mRNA | 37595498 | 37602288 | 100.0 | ID=Qrob_T0532570.2 |
| Qrob_Chr01 | mRNA | 38691337 | 38694866 | 97.0 | ID=Qrob_T0184060.2 |
| Qrob_Chr01 | mRNA | 38697663 | 38699156 | 93.3 | ID=Qrob_T0184050.2 |
| Qrob_Chr01 | mRNA | 55066440 | 55068599 | 100.0 | ID=Qrob_T0611170.2 |
| Qrob_Chr02 | mRNA | 17419146 | 17420228 | 97.3 | ID=Qrob_T0454110.2 |
| Qrob_Chr02 | mRNA | 28382238 | 28397120 | 100.0 | ID=Qrob_T0299620.2 |
| Qrob_Chr02 | mRNA | 28400301 | 28401018 | 0.0 | ID=Qrob_T0299610.2 |
| Qrob_Chr02 | mRNA | 28404084 | 28407207 | 100.0 | ID=Qrob_T0299600.2 |
| Qrob_Chr02 | mRNA | 28987060 | 28998888 | 99.0 | ID=Qrob_T0089890.2 |
| Qrob_Chr02 | mRNA | 29002128 | 29005412 | 100.0 | ID=Qrob_T0089880.2 |
| Qrob_Chr02 | mRNA | 34712751 | 34715369 | 94.4 | ID=Qrob_T0437960.2 |
| Qrob_Chr02 | mRNA | 42655612 | 42657345 | 100.0 | ID=Qrob_T0589320.2 |
| Qrob_Chr02 | mRNA | 44205331 | 44205970 | 100.0 | ID=Qrob_T0377200.2 |
| Qrob_Chr02 | mRNA | 44213324 | 44216710 | 99.0 | ID=Qrob_T0377190.2 |
| Qrob_Chr02 | mRNA | 46990675 | 46991783 | 100.0 | ID=Qrob_T0609890.2 |
| Qrob_Chr02 | mRNA | 46996815 | 47001368 | 100.0 | ID=Qrob_T0609880.2 |

| | | | | | |
|---|---|---|---|---|---|
| Qrob_Chr02 | mRNA | 47003403 | 47004571 | 100.0 | ID=Qrob_T0609870.2 |
| Qrob_Chr02 | mRNA | 49365511 | 49368425 | 100.0 | ID=Qrob_T0709860.2 |
| Qrob_Chr02 | mRNA | 49383529 | 49390427 | 100.0 | ID=Qrob_T0709880.2 |
| Qrob_Chr02 | mRNA | 50880650 | 50885782 | 88.1 | ID=Qrob_T0528660.2 |
| Qrob_Chr02 | mRNA | 50889161 | 50893177 | 100.0 | ID=Qrob_T0528650.2 |
| Qrob_Chr02 | mRNA | 50904986 | 50905468 | 0.0 | ID=Qrob_T0528630.2 |
| Qrob_Chr02 | mRNA | 53983164 | 53985503 | 100.0 | ID=Qrob_T0700420.2 |
| Qrob_Chr02 | mRNA | 53992359 | 54004961 | 95.0 | ID=Qrob_T0700440.2 |
| Qrob_Chr02 | mRNA | 92431869 | 92436245 | 97.0 | ID=Qrob_T0282290.2 |
| Qrob_Chr03 | mRNA | 40167291 | 40167596 | 100.0 | ID=Qrob_T0745850.2 |
| Qrob_Chr03 | mRNA | 40169165 | 40169425 | 0.0 | ID=Qrob_T0745840.2 |
| Qrob_Chr03 | mRNA | 40171988 | 40176838 | 100.0 | ID=Qrob_T0745830.2 |
| Qrob_Chr04 | mRNA | 27997801 | 27998563 | 90.1 | ID=Qrob_T0399160.2 |
| Qrob_Chr04 | mRNA | 41651662 | 41655844 | 0.0 | ID=Qrob_T0308170.2 |
| Qrob_Chr04 | mRNA | 41665218 | 41666719 | 100.0 | ID=Qrob_T0308190.2 |
| Qrob_Chr05 | mRNA | 16208833 | 16211432 | 100.0 | ID=Qrob_T0178490.2 |
| Qrob_Chr05 | mRNA | 17212822 | 17227633 | 98.0 | ID=Qrob_T0178670.2 |
| Qrob_Chr05 | mRNA | 17232249 | 17239413 | 100.0 | ID=Qrob_T0178680.2 |
| Qrob_Chr05 | mRNA | 2032128 | 2035537 | 91.1 | ID=Qrob_T0649050.2 |
| Qrob_Chr05 | mRNA | 2041278 | 2044530 | 0.0 | ID=Qrob_T0649030.2 |
| Qrob_Chr05 | mRNA | 22675622 | 22677555 | 39.1 | ID=Qrob_T0583160.2 |
| Qrob_Chr05 | mRNA | 22684391 | 22686918 | 100.0 | ID=Qrob_T0583150.2 |
| Qrob_Chr05 | mRNA | 22704507 | 22708866 | 100.0 | ID=Qrob_T0013730.2 |
| Qrob_Chr05 | mRNA | 22714427 | 22722619 | 95.0 | ID=Qrob_T0013740.2 |
| Qrob_Chr05 | mRNA | 23925445 | 23929495 | 22.3 | ID=Qrob_T0523430.2 |
| Qrob_Chr05 | mRNA | 23941803 | 23946877 | 100.0 | ID=Qrob_T0523410.2 |
| Qrob_Chr05 | mRNA | 28012467 | 28013501 | 0.0 | ID=Qrob_T0449720.2 |

| | | | | | |
|---|---|---|---|---|---|
| Qrob_Chr05 | mRNA | 7906041 | 7912235 | 100.1 | ID=Qrob_T0683210.2 |
| Qrob_Chr06 | mRNA | 21260908 | 21264170 | 54.0 | ID=Qrob_T0005740.2 |
| Qrob_Chr06 | mRNA | 27193253 | 27195122 | 98.0 | ID=Qrob_T0256040.2 |
| Qrob_Chr06 | mRNA | 28622710 | 28630237 | 100.1 | ID=Qrob_T0747960.2 |
| Qrob_Chr06 | mRNA | 40963979 | 40968076 | 100.0 | ID=Qrob_T0413630.2 |
| Qrob_Chr06 | mRNA | 40974308 | 40977654 | 0.0 | ID=Qrob_T0413610.2 |
| Qrob_Chr07 | mRNA | 37924676 | 37927169 | 1.5 | ID=Qrob_T0131310.2 |
| Qrob_Chr07 | mRNA | 37930043 | 37933882 | 92.1 | ID=Qrob_T0131320.2 |
| Qrob_Chr07 | mRNA | 4037164 | 4038276 | 95.1 | ID=Qrob_T0379860.2 |
| Qrob_Chr07 | mRNA | 4038746 | 4042140 | 37.5 | ID=Qrob_T0379870.2 |
| Qrob_Chr07 | mRNA | 41623546 | 41636343 | 100.0 | ID=Qrob_T0090900.2 |
| Qrob_Chr07 | mRNA | 46049058 | 46056801 | 92.1 | ID=Qrob_T0407530.2 |
| Qrob_Chr07 | mRNA | 7062267 | 7067123 | 20.0 | ID=Qrob_T0265770.2 |
| Qrob_Chr08 | mRNA | 10656567 | 10658485 | 100.0 | ID=Qrob_T0132290.2 |
| Qrob_Chr08 | mRNA | 25970299 | 25973975 | 100.0 | ID=Qrob_T0441200.2 |
| Qrob_Chr08 | mRNA | 25978580 | 25979250 | 100.0 | ID=Qrob_T0441190.2 |
| Qrob_Chr08 | mRNA | 50962263 | 50985338 | 92.2 | ID=Qrob_T0437680.2 |
| Qrob_Chr08 | mRNA | 52158757 | 52169697 | 100.0 | ID=Qrob_T0631150.2 |
| Qrob_Chr08 | mRNA | 52173071 | 52176597 | 100.0 | ID=Qrob_T0631140.2 |
| Qrob_Chr08 | mRNA | 52181642 | 52185377 | 100.0 | ID=Qrob_T0631130.2 |
| Qrob_Chr08 | mRNA | 62286008 | 62293369 | 93.0 | ID=Qrob_T0413010.2 |
| Qrob_Chr08 | mRNA | 62295320 | 62296804 | 100.0 | ID=Qrob_T0413000.2 |
| Qrob_Chr08 | mRNA | 62298760 | 62300062 | 100.0 | ID=Qrob_T0412990.2 |
| Qrob_Chr08 | mRNA | 62746427 | 62753205 | 100.0 | ID=Qrob_T0605490.2 |
| Qrob_Chr08 | mRNA | 62756677 | 62757416 | 100.0 | ID=Qrob_T0605480.2 |
| Qrob_Chr08 | mRNA | 62783709 | 62786511 | 100.0 | ID=Qrob_T0605470.2 |
| Qrob_Chr08 | mRNA | 62794403 | 62797526 | 99.0 | ID=Qrob_T0211590.2 |

| | | | | | |
|---|---|---|---|---|---|
| Qrob_Chr09 | mRNA | 26419438 | 26420323 | 100.0 | ID=Qrob_T0285940.2 |
| Qrob_Chr09 | mRNA | 26422819 | 26423321 | 92.1 | ID=Qrob_T0285930.2 |
| Qrob_Chr09 | mRNA | 41156949 | 41176337 | 98.0 | ID=Qrob_T0489020.2 |
| Qrob_Chr10 | mRNA | 48860710 | 48861429 | 0.0 | ID=Qrob_T0644820.2 |
| Qrob_Chr10 | mRNA | 48864519 | 48865241 | 0.0 | ID=Qrob_T0644810.2 |
| Qrob_Chr11 | mRNA | 15021331 | 15022690 | 25.0 | ID=Qrob_T0104460.2 |
| Qrob_Chr11 | mRNA | 33429757 | 33451483 | 100.0 | ID=Qrob_T0010570.2 |
| Qrob_Chr11 | mRNA | 51079119 | 51086901 | 100.0 | ID=Qrob_T0251500.2 |
| Qrob_Chr11 | mRNA | 51094139 | 51094775 | 0.0 | ID=Qrob_T0251490.2 |
| Qrob_Chr12 | mRNA | 32347782 | 32354615 | 99.0 | ID=Qrob_T0149640.2 |

**Table S2.** Summary table of Q. robur candidate genes under recent selection detected by both SweeD and OmegaPlus ($p < .01$).

| **ID** | **Source** | **Type** | **Start** | **End** | **Score** | **Strand** | **Gene ID** |
|---|---|---|---|---|---|---|---|
| Qrob_Chr01 | Egn | mRNA | 21470613 | 21471928 | 98.7 | + | ID=Qrob_T0000290.2 |
| Qrob_Chr01 | egn | mRNA | 47849967 | 47852265 | 99 | + | ID=Qrob_T0041720.2 |
| Qrob_Chr02 | egn | mRNA | 9550828 | 9552258 | 0 | - | ID=Qrob_T0433250.2 |
| Qrob_Chr03 | egn | mRNA | 36770831 | 36774503 | 100 | + | ID=Qrob_T0102900.2 |
| Qrob_Chr03 | egn | mRNA | 49666659 | 49673966 | 27.1 | - | ID=Qrob_T0170450.2 |

| | | | | | | | |
|---|---|---|---|---|---|---|---|
| Qrob_Chr09 | egn | mRNA | 13945795 | 13946619 | 0 | + | ID=Qrob_T0542880.2 |
| Qrob_Chr11 | egn | mRNA | 5403212 | 5411610 | 100 | + | ID=Qrob_T0080510.2 |
| Qrob_Chr11 | Manual_v2 | mRNA | 44537895 | 44539098 | . | - | Name=Qrob_T0158900.2 |
| Qrob_Chr11 | egn | mRNA | 38960405 | 38963858 | 98 | + | ID=Qrob_T0699490.2 |
| Qrob_Chr12 | egn | mRNA | 33369924 | 33377321 | 8.1 | + | ID=Qrob_T0662860.2 |

**Table S3.** Length variants in base pairs and point mutations detected in cpDNA fragments of five identified haplotypes. Key features used to match haplotypes are underlined. A) DT fragment digested with TaqI. B) AS fragment digested with HinfI. C) CD fragment digested with TaqI. D) TF fragment digested with AluI. E) Point mutations in the DT and TF fragments, identified with AluI and CfoI, respectively.

**A.**

| **Haplotypes (this study)** | **Haplotypes (Petit et al., 2002b)** | **Lineage** | **DT1** | **DT2** | **DT3** | **DT4** |
|---|---|---|---|---|---|---|
| I | 10 | B | 571 | 389 | <u>288</u> | 215 |
| II | 11 | B | 571 | 389 | <u>288</u> | 215 |
| III, IV | 12 | B | 571 | 389 | <u>287</u> | 215 |
| V | 7,26 | A | 571 | 389 | <u>215</u> | 211 |

**B.**

| **Haplotypes (this study)** | **Haplotypes (Petit et al., 2002b)** | **Lineage** | **AS1** | **AS2** | **AS3** | **AS4** | **AS5** | **AS6** |
|---|---|---|---|---|---|---|---|---|
| I | 10 | B | 677 | 569 | 516 | 370 | 310 | 211 |
| II | 11 | B | 677 | 569 | 516 | 370 | 310 | 210 |
| III, IV | 12 | B | 677 | 569 | 516 | 370 | 310 | 211 |
| V | 7,26 | A | <u>649</u> | 569 | 516 | 370 | 310 | 210 |

**C.**

| **Haplotypes (this study)** | **Haplotypes (Petit et al., 2002b)** | **Lineage** | **CD1** | **CD2** | **CD3** | **CD4** | **CD5** | **CD6** |
|---|---|---|---|---|---|---|---|---|
| I | 10 | B | 972 | 658 | 533 | 322 | 291 | 259 |
| II | 11 | B | 972 | 658 | 533 | 322 | 291 | 259 |
| III, IV | 12 | B | 972 | 658 | 533 | 322 | 291 | 259 |
| V | 7,26 | A | 972 | 658 | 533 | 322 | 291 | 259 |

**D.**

| **Haplotypes (this study)** | **Haplotypes (Petit et al., 2002b)** | **Lineage** | **TF1** | **TF2** | **TF3** | **TF4** | **TF5** | **TF6** |
|---|---|---|---|---|---|---|---|---|
| I | 10 | B | 981 | 665 | 89 | 65 | 45 | 30 |
| II | 11 | B | 981 | 678 | 89 | 65 | 45 | 30 |
| III, IV | 12 | B | 981 | 665 | 89 | 65 | 45 | 30 |
| V | 7,26 | A | 980 | 664 | 89 | 65 | 45 | 30 |

**E.**

| **Haplotypes (this study)** | **Haplotypes (Petit et al., 2002b)** | **Lineage** | **DT-AluI** | **TF-CfoI** |
|---|---|---|---|---|
| I | 10 | B | Yes | No |
| II | 11 | B | Yes | No |
| III, IV | 12 | B | Yes | No |
| V | 7, 26 | A | No | Yes |

**Figure S1.** PCA of 386 oak trees of two species based on 2,768,547 SNPs ($r^2 < 0.4$). Colours represent sites while symbols represent species. A) PC1 against PC2. B) PC1 against PC3. C) PC2 against PC3. D) Eigenvalues of the computed principal components.

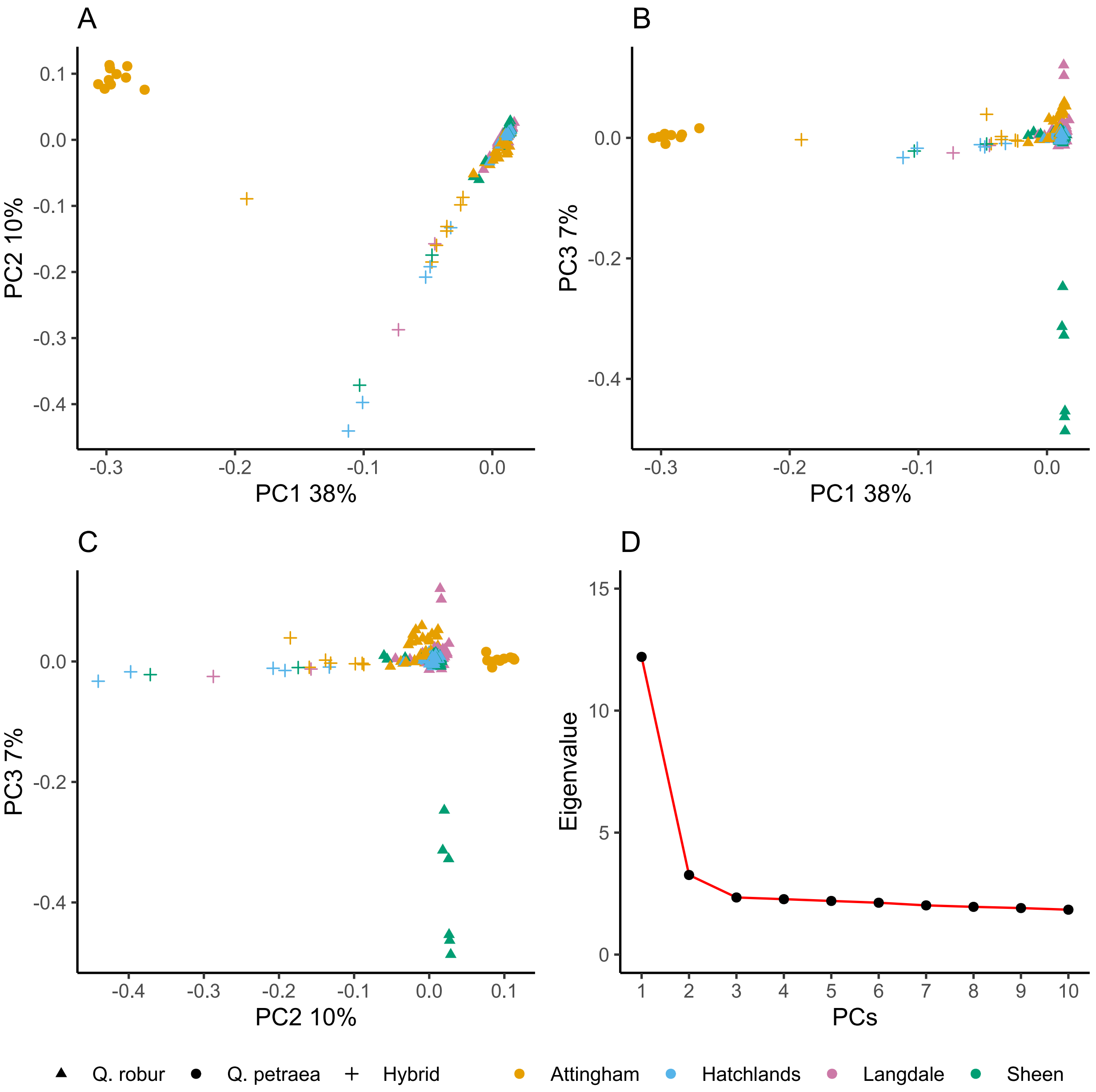

**Figure S2.** Genome-wide, genic and intergenic interspecific $F_{st}$ distribution between *Q. robur* and *Q. petraea* based on 914,242, 223,319 and 690,923 SNPs, respectively. Genome-wide mean, median and standard deviation: 0.158, 0.111 and 0.156. Genic regions mean, median and standard deviation: 0.155, 0.111 and 0.153. Intergenic regions mean, median and standard deviation: 0.159, 0.111 and 0.157.

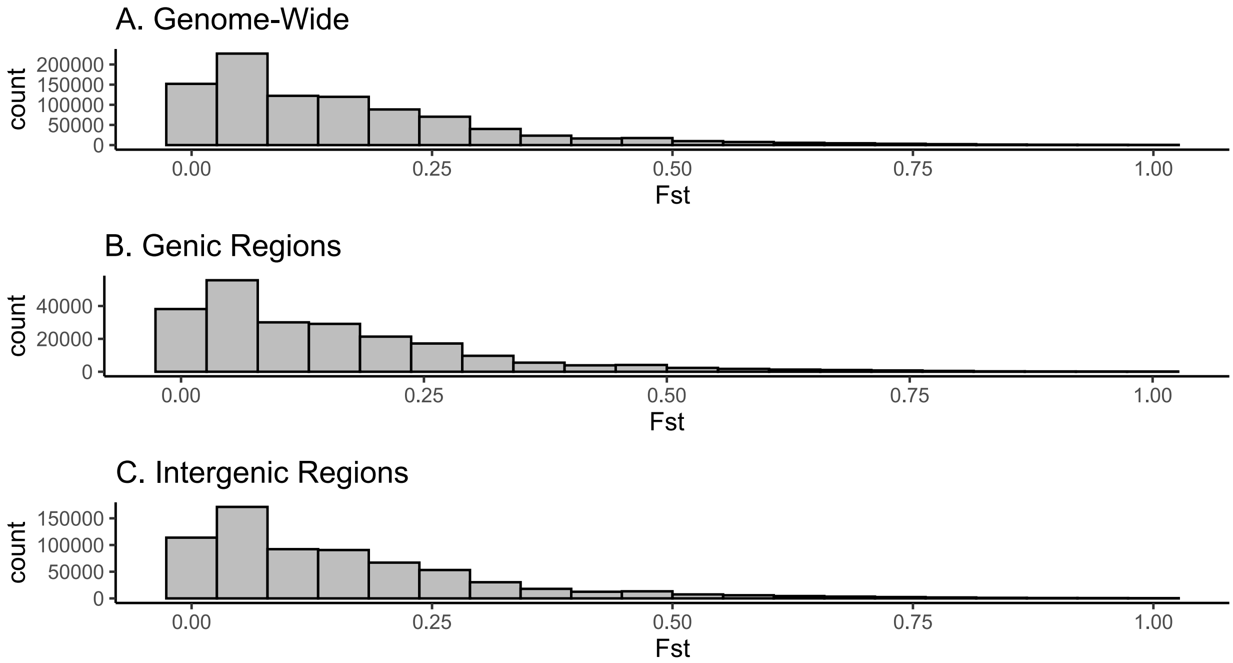

**Figure S3.** $F_{st}$/Heterozygosity distribution computed with the R package “fsthet”, based on 914,242 genome-wide SNPs between 10 *Q. robur* and 10 *Q. petraea* individuals. Red lines delimit the 0.99 confidence envelope. Points with $F_{st}$ > 0.6 and outside this envelope were considered outliers.

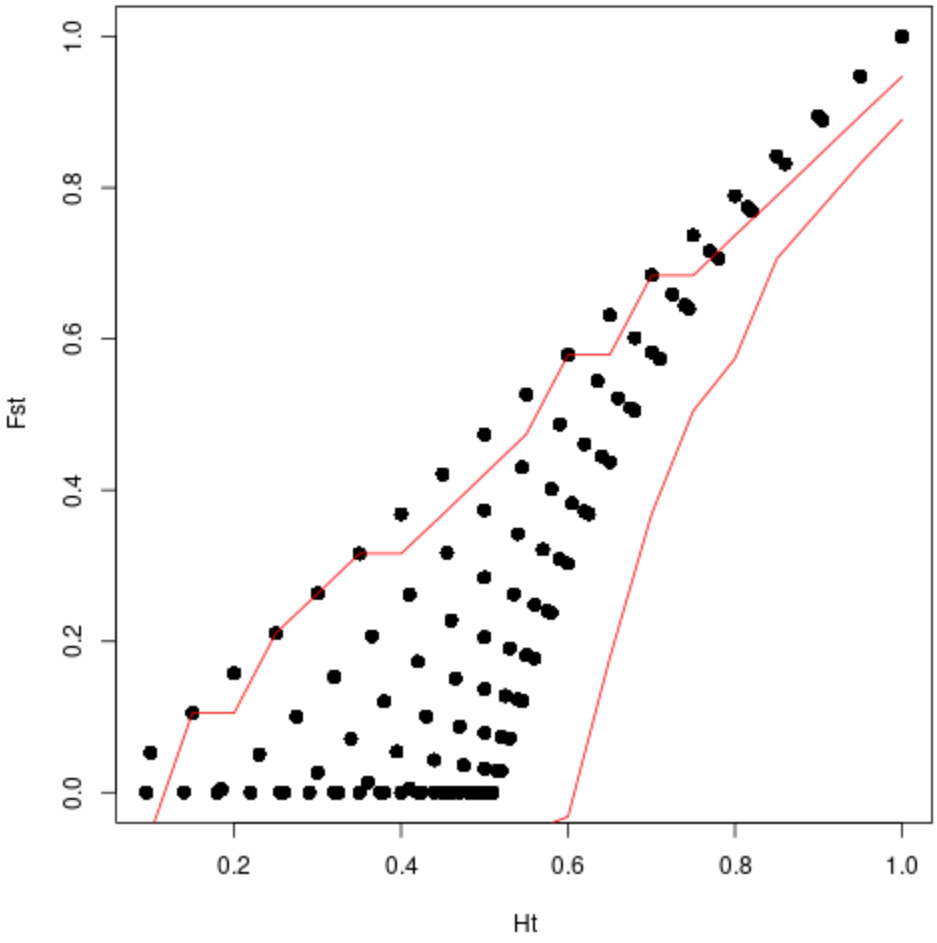

**Figure S4.** PCA of 360 *Q. robur* individual based on 839,911 unlinked SNPs ($r^2 < 0.4$). Colours represent sites. A) PC1 against PC2. B) PC1 against PC3. C) PC2 against PC3. D) Eigenvalues of the computed principal components.

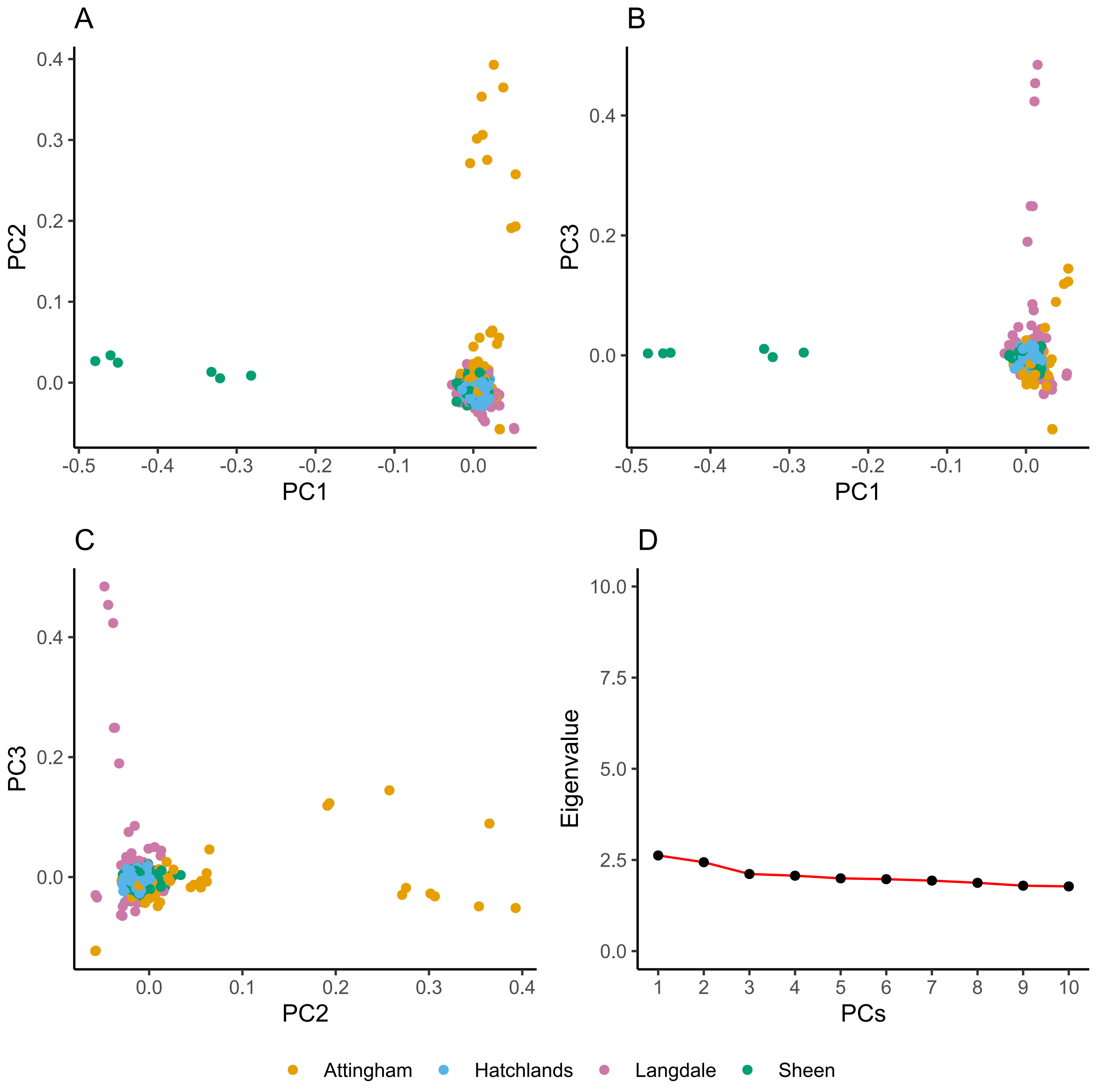

**Figure S5.** Distribution of genomic relatedness (kin) between 360 *Q. robur* trees based on 839,911 unlinked ($r^2 < 0.4$) SNPs with minor allele frequencies above 0.05.

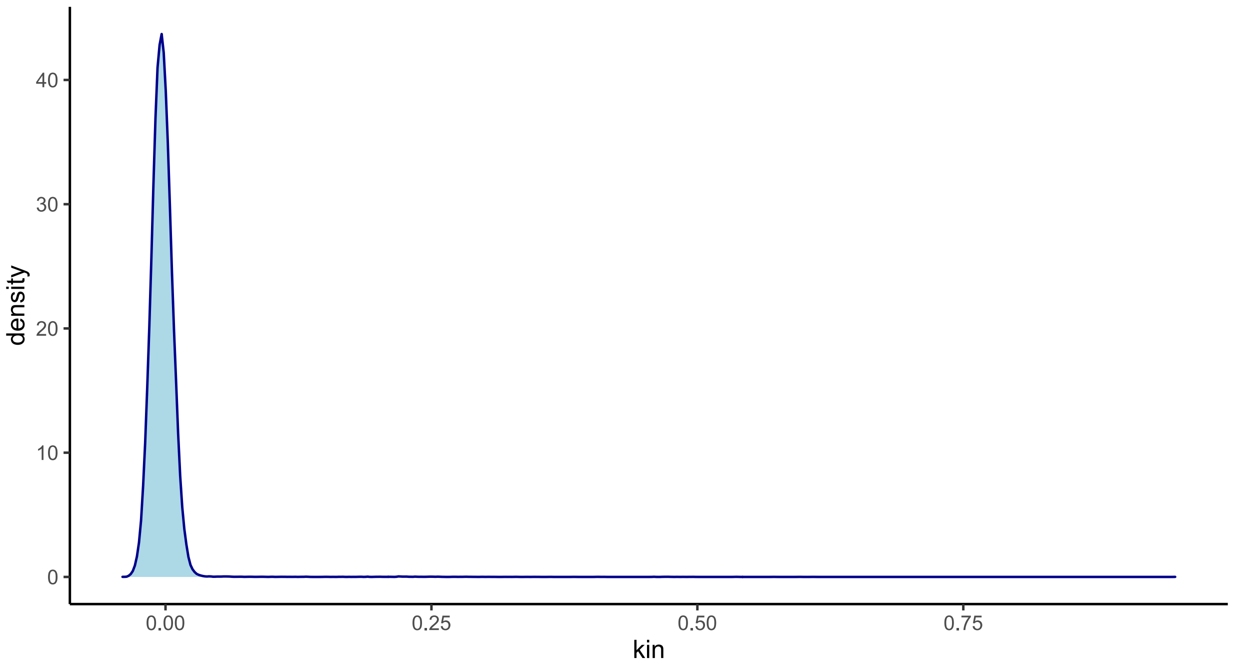

**Figure S6.** Marker based genomic relatedness within parkland sites, calculated according to the formulas in vanRaden (2008). Diagonals represent self-to-self relatedness.

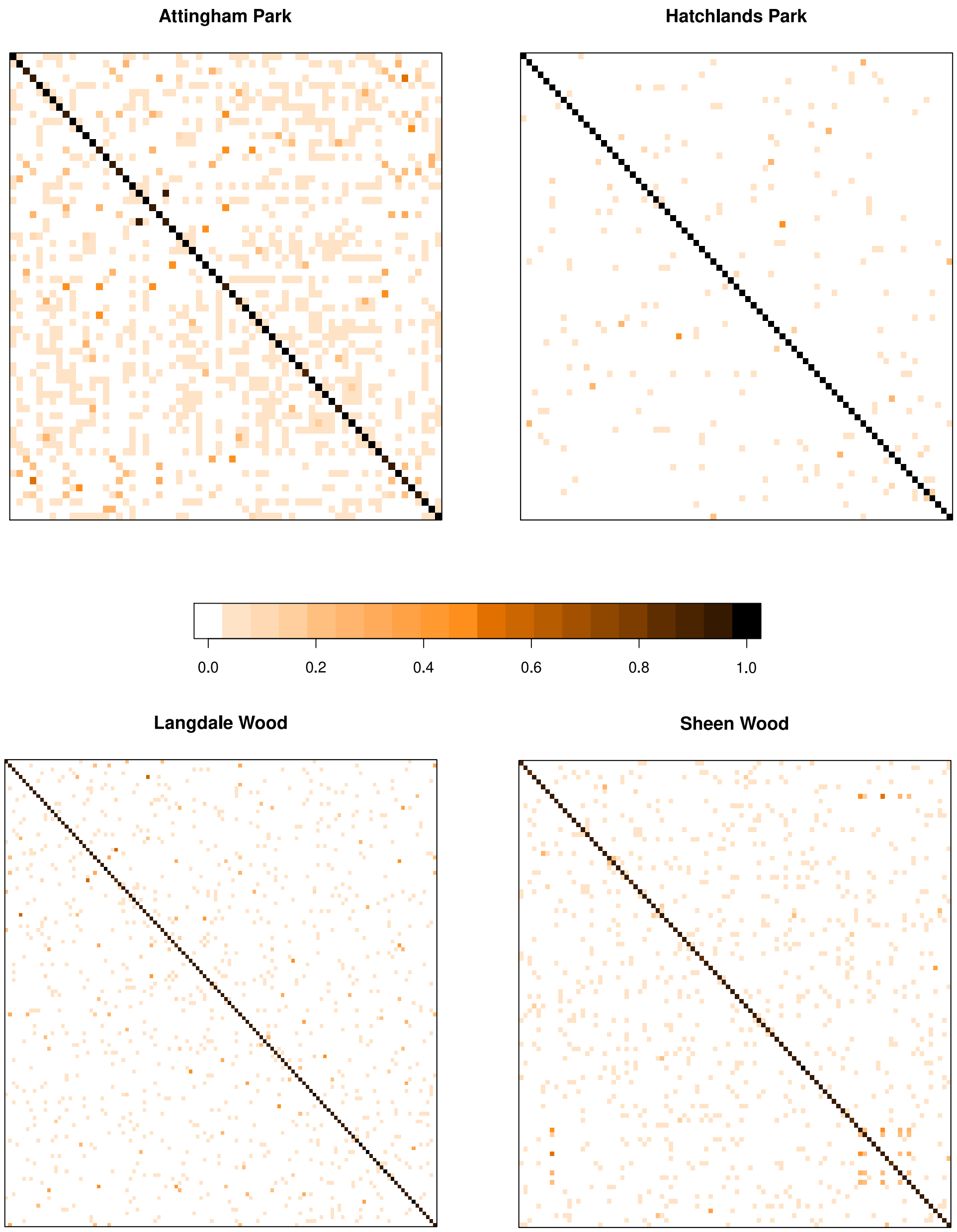

**Figure S7.** PCA of 261 unrelated (kin < 0.05) *Q. robur* individuals based on 839,911 SNPs (r2 < 0.4) . Colours represent sites. A) PC1 against PC2. B) PC1 against PC3. C) PC2 against PC3. D) Plot of eigenvalues for the computed principal components.

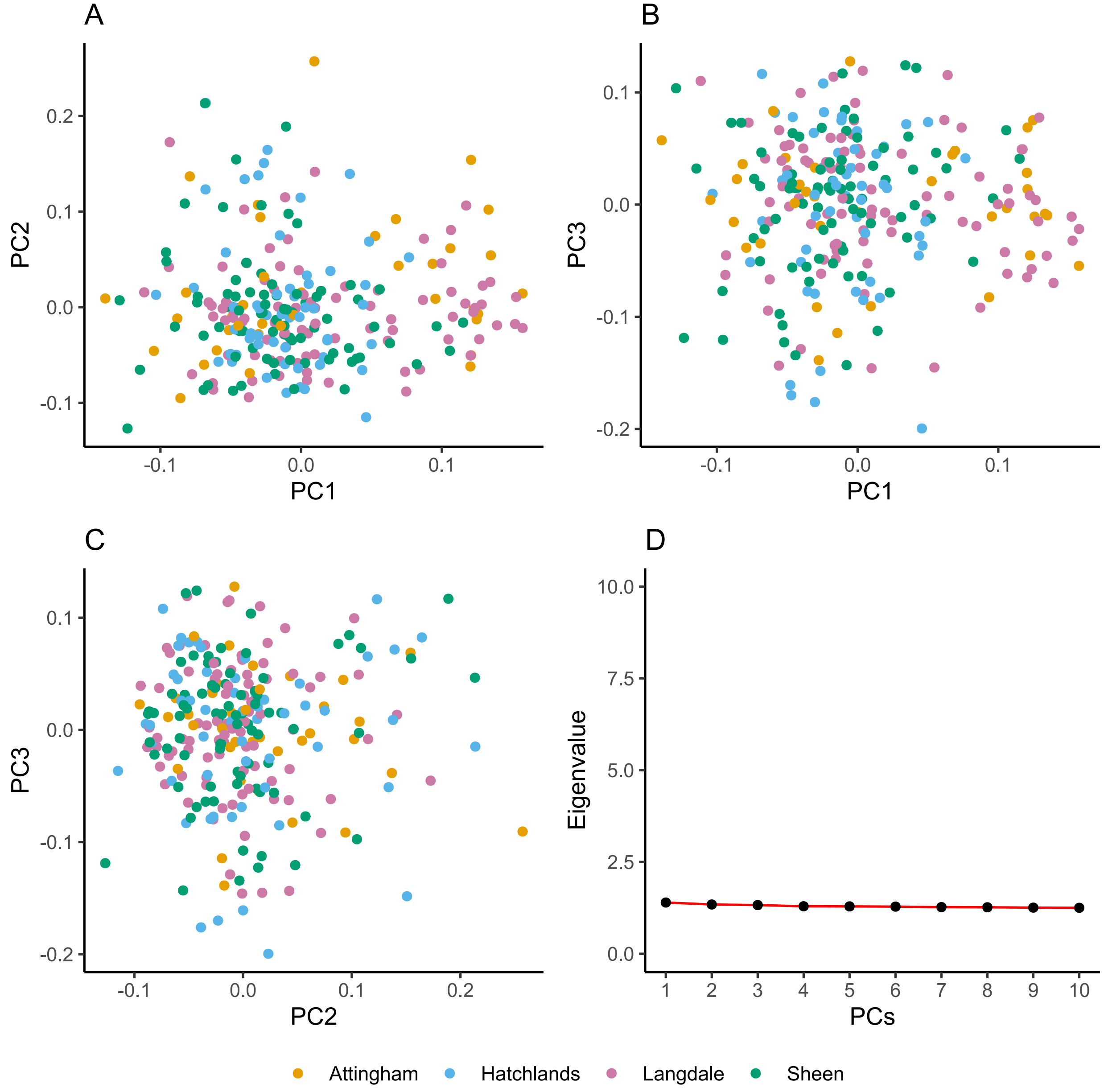

**Figure S8.** Linkage disequilibrium (LD) decay in the *Q. robur* genome. A) LD decay, estimated with Plink $r^2$ function. Points are 500 bases apart. B) Linkage disequilibrium block size distribution.

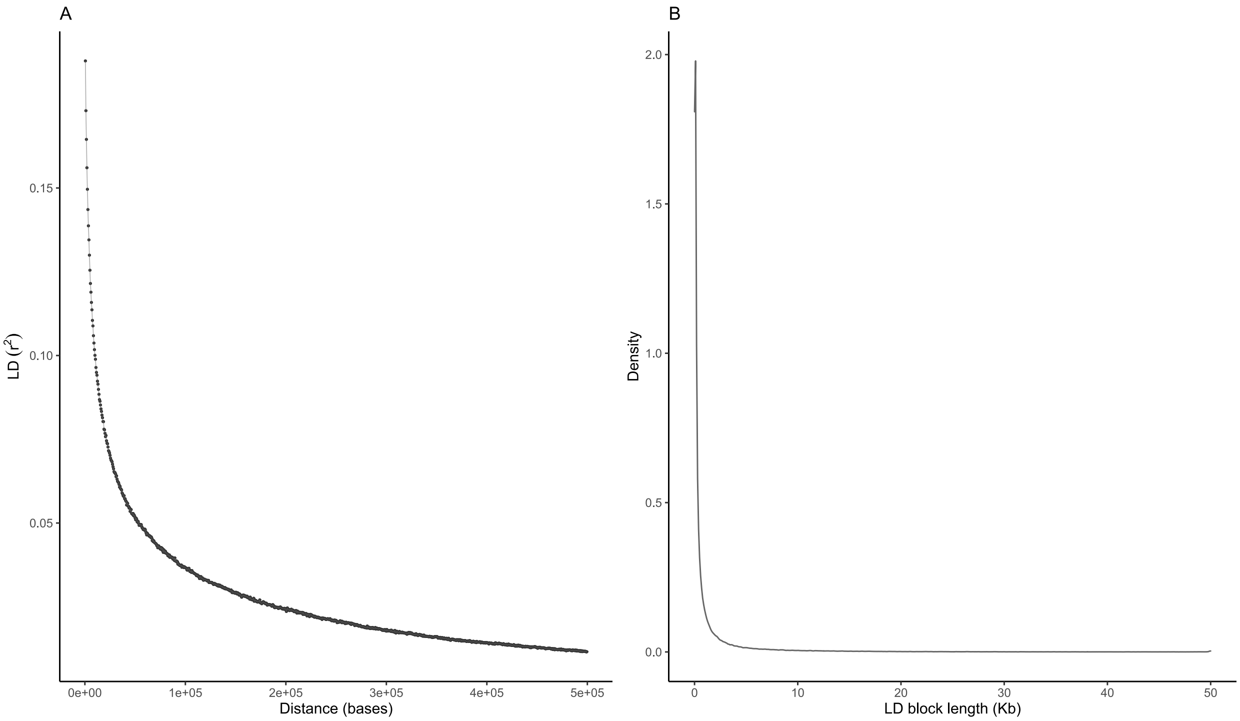

**Figure S9.** Selective sweep scan output for chromosomes with common outliers between SweeD and OmegaPlus. (AB) The x axis denotes the base pair position on the chromosome, and the y axis shows the CLR and ω statistic computed with SweeD and OmegaPlus, respectively. (C) Combined plot for SweeD and OmegaPlus. Red dots represent outliers in common between SweeD and OmegaPlus ($p < .01$).

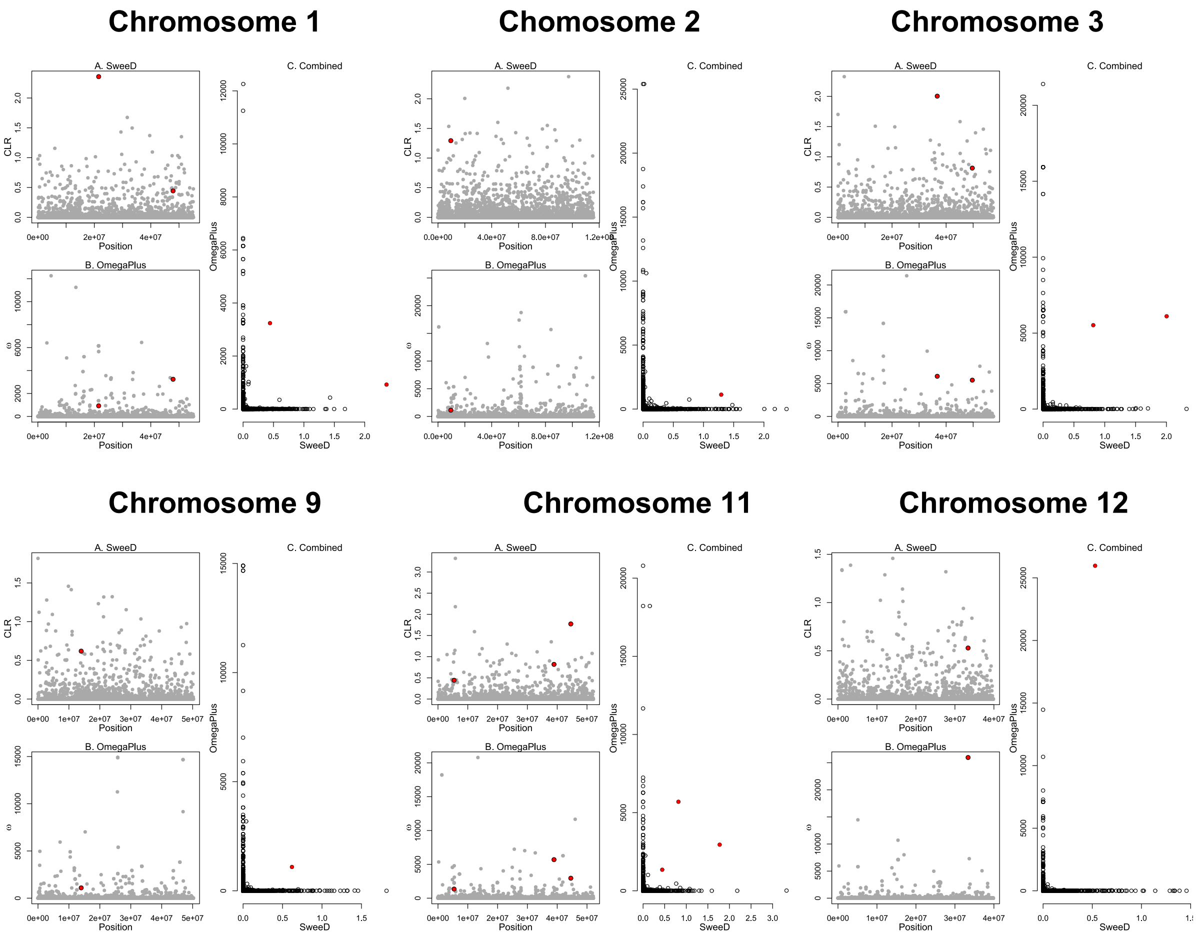

**Figure S10.** Distribution of the four chloroplast haplotypes identified between sampled parkland sites and species.

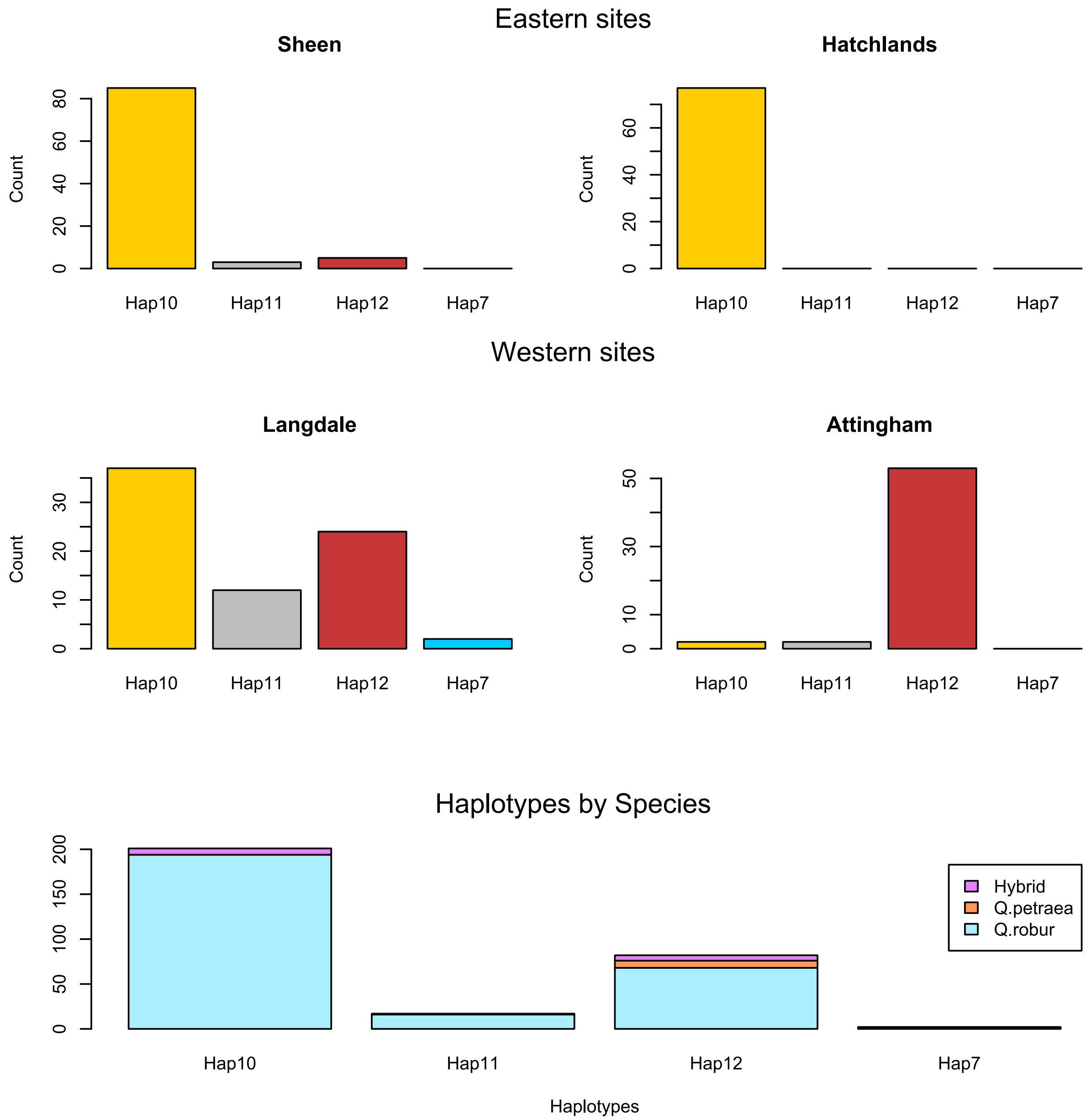

**Figure S11.** Haplotypes frequency contour regions estimated with ordinary kriging interpolation of the chloroplast data of 178 ancient woodlands (Cottrell et al., 2002). Yellow: > 50%. Orange: 60-70%. Red: > 70%. Black points represent the sites sampled in this study.

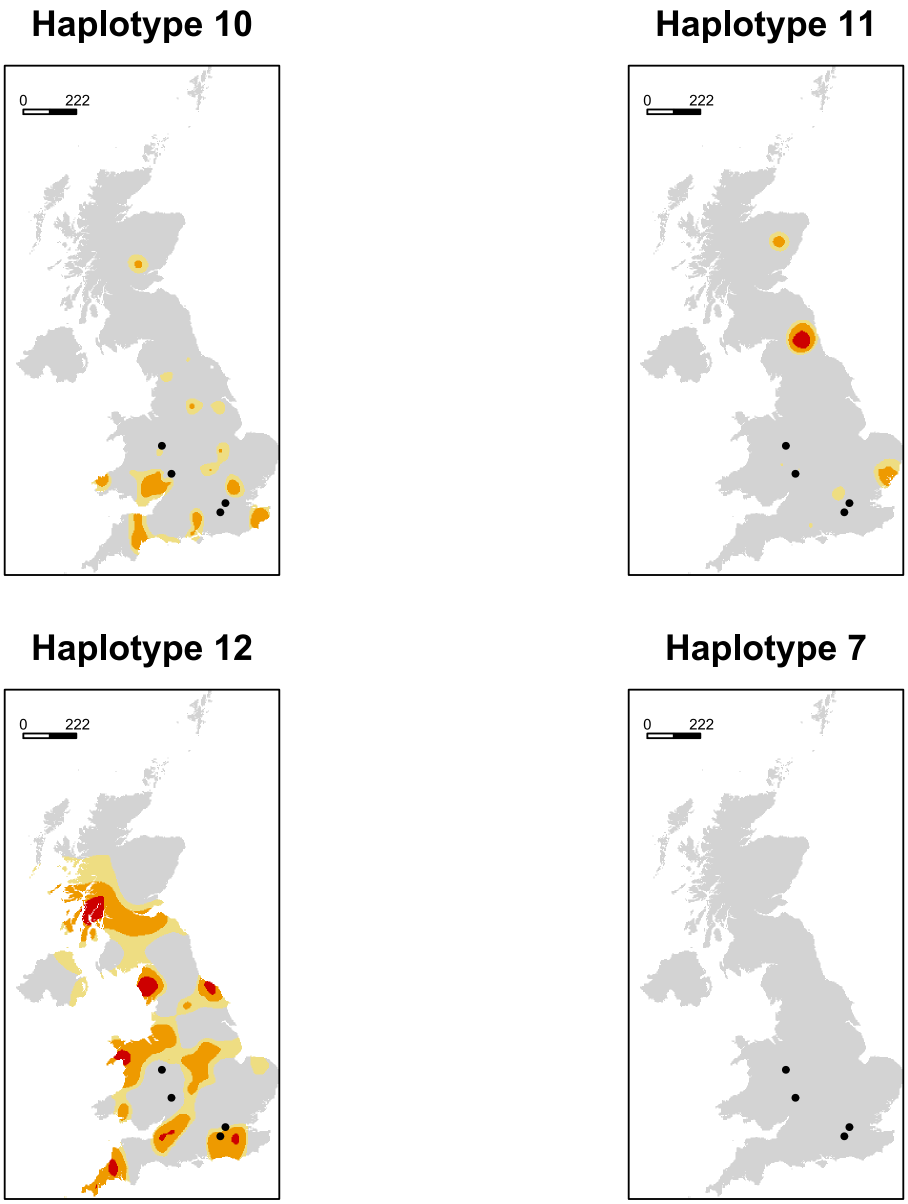
